# Mitochondria are absent from microglial processes performing surveillance, chemotaxis, and phagocytic engulfment

**DOI:** 10.1101/2024.10.15.618505

**Authors:** Alicia N. Pietramale, Xhoela Bame, Megan E. Doty, Robert A. Hill

## Abstract

Microglia continually surveil the brain allowing for rapid detection of tissue damage or infection. Microglial metabolism is linked to tissue homeostasis, yet how mitochondria are subcellularly partitioned in microglia and dynamically reorganize during surveillance, injury responses, and phagocytic engulfment in the intact brain are not known. Here, we performed intravital imaging of microglia mitochondria, revealing that microglial processes diverge, with some containing multiple mitochondria while others are completely void. Microglial processes that engage in minute-to-minute surveillance typically do not have mitochondria. Moreover, unlike process surveillance, mitochondrial motility does not change with animal anesthesia. Likewise, the processes that acutely chemoattract to a lesion site or initially engage with a neuron undergoing programmed cell death do not contain mitochondria. Rather, microglia mitochondria have a delayed arrival into the responding cell processes. Thus, there is subcellular heterogeneity of mitochondrial partitioning and asymmetry between mitochondrial localization and cell process motility or acute damage responses.

## INTRODUCTION

Microglia are the most dynamic cell in the central nervous system^1,2^. As the resident innate immune cell, their functions span tissue surveillance to pathological responses. They do this by continually extending and retracting their highly branched processes to probe the environment for infection or injury. In the case of a focal injury, microglia display immediate process chemotaxis, effectively barricading the injury site to prevent further damage to the parenchyma^3,4^. Subsequent microglial phagocytosis mediates engulfment and clearance of cellular debris, ultimately facilitating a return to homeostasis from this inflammatory injury. Microglia are also capable of performing phagocytosis without inflammatory stimuli via specialized phagocytic receptors that recognize damaged or dying cells or cell compartments^5–9^. Together, surveillance, chemotaxis, and phagocytosis involve radical morphological changes in microglia, but our understanding of how the internal cellular architecture changes alongside these cell behaviors is incomplete.

Mitochondria are signaling hubs involved in many processes ranging from cell metabolism, calcium handling, cell death and proliferation, and the generation of reactive oxygen species (ROS)^10,11^. Mitochondrial subcellular localization influences these multifunctional roles and their proximity to other organelles or positioning near cell-cell contacts enhances their ability to integrate and respond to intracellular and extracellular stimuli^10^. Mitochondria are highly dynamic organelles capable of transport throughout the cell mediated by the cytoskeleton, and changing shape through fission and fusion events, each of which have been linked to the metabolic state of the cell^10,12–14^.

In aging and neurodegeneration microglial function is dysregulated, potentially contributing to the pathology rather than helping the system return to homeostasis^2,15–24^. In this state, microglia exhibit mitochondrial dysregulation shown by increased ROS emission, mitochondrial DNA damage, and metabolic dysfunction^24–27^. Additionally, inflammatory environments alter mitochondrial shape in microglia in vitro, which coincides with a microglial bioenergetic shift and an increase in pro-inflammatory cytokine release^28–31^. Moreover, targeted manipulation of distinct mitochondrial genes within microglia has revealed diverse roles for mitochondria in microglial physiology and pathology^32–36^. However, it remains to be explored how mitochondrial shape and motility change as microglia perform surveillance, phagocytosis, and respond to injury in the healthy brain.

In this study, we define the microglia mitochondrial dynamics at homeostasis to provide a baseline of the mitochondrial phenotype as microglia carry out their multifaceted functions. To investigate microglia mitochondria, we generated a triple transgenic mouse with microglia and their mitochondria endogenously labeled with separate fluorescent markers. This enables us to study mitochondrial morphology and motility in live animals while maintaining microglia in their native microenvironment. Together our data establish the dynamics and morphometrics of microglia mitochondria in vivo and indicate heterogeneity of process function within a single microglial cell.

## RESULTS

### Mitochondrial morphometrics in mouse neocortical microglia

Inflammatory stimuli shift the bioenergetics of cultured microglia which coincides with a morphological change of mitochondria in microglia^28–31^. However, mitochondrial morphometrics (content, shape, and subcellular distribution) and motility have not been characterized in microglia in the healthy, live, and intact brain. To visualize mitochondria specifically within microglia we generated a triple transgenic mouse line, *Cx3cr1*-creER: Ai9: PhAM, enabling dual fluorophore expression in the cell cytoplasm (Ai9) and mitochondrial matrix (Dendra2) driven by tamoxifen inducible cre recombination (Fig. 1a, Supplementary Fig. 1a, b, and Supplementary Movie 1). To make the brain optically accessible for intravital imaging, cranial window surgeries were performed over the somatosensory cortex (Supplementary Fig. 1a). Immediately after the surgery, high resolution optical images were taken, then the microglia cytoplasm and mitochondria were 3 dimensionally (3D) segmented (Supplementary Fig. 1c). Using the volumes extracted from the 3D segmentations, we found the mitochondrial to cytoplasmic volume fraction in the whole cell and the compartments: soma and processes to be respectively ∼10%, 17%, and ∼8% (Fig. 1b). The soma contained relatively few but large mitochondrial networks by volume while the mitochondria occupying the processes were smaller in volume but greater in number (Fig. 1a, c). Mitochondrial shape was also determined within these cellular compartments with circularity and aspect ratio measurements as mitochondrial morphology has been correlated to metabolic fitness^13,37^. The circularity of the soma and process mitochondria showed no difference; however, the aspect ratio showed that the process mitochondria were more elongated than the soma mitochondria (Fig. 1f).

**Figure 1:**
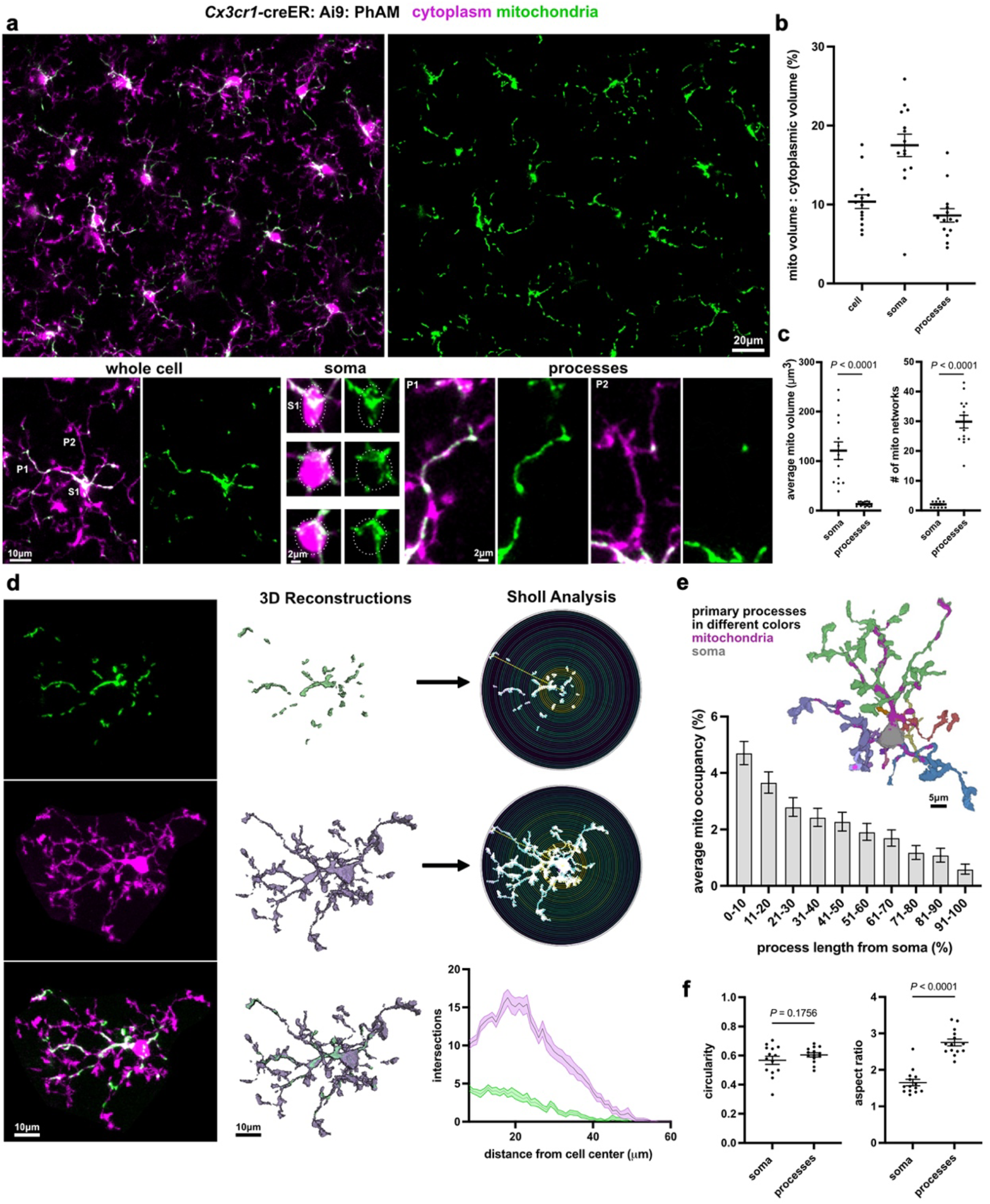
Mitochondrial morphometrics in mouse neocortical microglia. **a)** In vivo image of mitochondrial labeling (green) in microglia (magenta) in the upper layers of the mouse somatosensory cortex. The images on the bottom show mitochondrial partitioning in 1 microglia (left image), representative soma from three separate microglia (middle images) and the two processes cropped from the first microglia as indicated and numbered. **b)** The mitochondrial to cytoplasm volume fraction for a subset of microglia that were segmented in 3D. Values are shown for the total cell, the soma, and the processes. Each dot in the graph represents 1 cell. **c)** Differences in the average mitochondrial network volume and number of mitochondrial networks between the soma and processes. **d)** 3D segmentation of a single microglia and its mitochondria and morphological analysis via Sholl reveals increased process complexity (magenta line) at 20 microns from the cell center but linear mitochondrial signals (green line) radiating from the center of the cell. **e)** The mitochondrial occupancy along the length of a microglial primary process in 10% bins starting from the center of the cell (0-10%) and going to the tip of the process (91-100%) (n = 117 primary processes from 14 cells from 6 mice, error bars indicate s.e.m.) **f)** Mitochondrial circularity and aspect ratio in the soma and processes. (for (**b**-**f**), n = 14 cells from 6 mice, paired two-tailed t-test, the line is at the mean ± s.e.m for error bars or error bands.)

To determine where mitochondria reside within microglia, sholl analysis was done on the cytoplasmic and mitochondrial segmentations. The microglia branching complexity increased as the density of branching increased (peaking at ∼20um away from the cell center) and then dropped when moving to the tips of the processes; however, this peak increase in complexity was not observed with the mitochondrial sholl analysis which linearly decreased when radiating away from the cell center to the tips of the processes (Fig. 1d). To measure this in a different way, the mitochondrial occupancy in individual primary processes was determined. A primary process was identified from the branch shooting off from the soma and all connected branches (see primary processes labeled in different colors in Fig. 1e). Like the sholl analysis, mitochondrial occupancy linearly decreased moving from the soma to the tips of the primary processes (Fig. 1e). Therefore, the complexity of the microglia cytoplasm does not dictate where there will be an increase in mitochondria presence. These analyses establish the baseline for mitochondrial morphometrics in the healthy intact brain and can serve as a reference as to how mitochondrial morphometrics may change in neuroinflammatory and neurodegenerative states.

### Mitochondrial morphometrics in human neocortical microglia

To determine if the microglial mitochondrial morphometrics were consistent across species we used a publicly available human electron microscopy dataset (H01) that was collected from 1 mm^3^ of the temporal cortex^38^. We selected 6 microglia to segment and confirmed their identity based on nuclear ultrastructure and cell cytoplasmic morphology^39^, then created 3D segmentations of the cytoplasm, nucleus, and mitochondria (Fig. 2a, Supplementary Fig. 2 and Supplementary Movie 2). The mitochondria to cytoplasmic volume fraction in the whole cell, soma, and processes was between ∼2.5% and ∼3.5% (Fig. 2a, d). We also found that the mitochondria in the soma were larger in volume but fewer in quantity, whereas the mitochondria in the processes were smaller in volume but greater in quantity (Fig. 2b, c, e). Moreover, the sholl analysis showed that mitochondria are more abundant at the cell center and linearly decrease to less abundance at the process tips (Fig. 2f). Not only are these findings consistent with the mouse mitochondrial morphometrics, but this electron microscopy dataset also provides the resolution to distinguish individual mitochondria and validates our findings made with the resolution capabilities of fluorescence microscopy.

**Figure 2:**
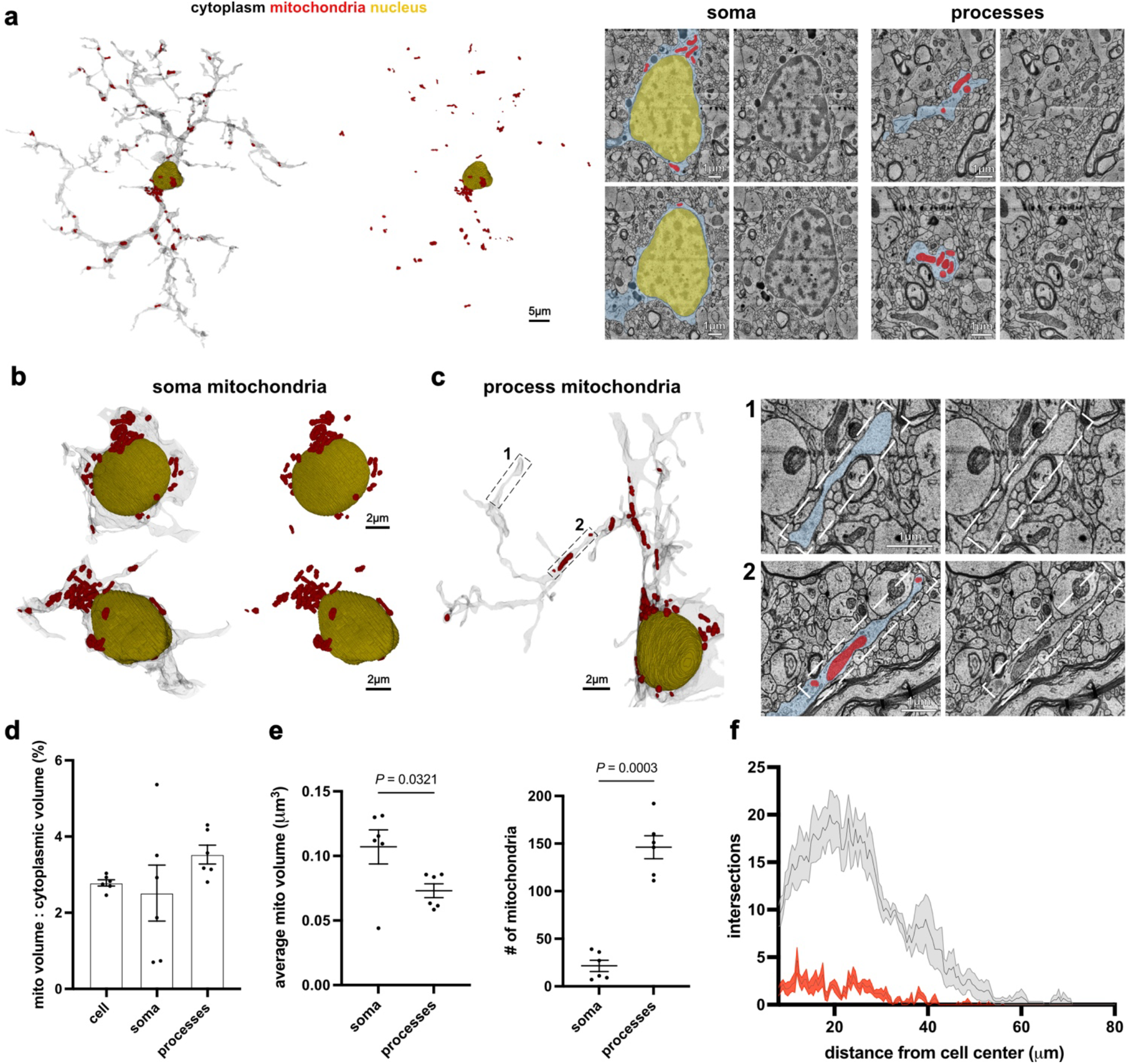
Mitochondrial morphometrics in human neocortical microglia. **a)** 3D reconstruction of a microglial cell (gray or blue), its nucleus (yellow), and all its mitochondria (red) taken from a sample of human neocortex. The images on the right show examples of electron micrographs taken through the soma or single processes that were used for cell and organelle segmentations. **b)** 3D renderings of microglial soma showing examples of perinuclear mitochondria. **c)** Some microglial processes contain many mitochondria and others are completely void as shown by the reconstruction (left) and the electron micrographs (right) of the indicated regions. **d)** The mitochondrial to cytoplasm volume fraction for the microglia that were segmented in 3D. Values are shown for the total cell, the soma, and the processes. Each dot in the graph represents 1 cell **e)** Differences in the total mitochondrial volume and number of mitochondrial networks between the soma and processes (paired two-tailed t-test). **f)** Sholl analysis reveals increased process complexity (gray line) at 20 microns from the cell center but linear mitochondrial signals (red line) measured from the center of the cell (for (**d-f**), n = 6 cells, error bars or bands indicate s.e.m.).

### Mitochondria are heterogeneously partitioned throughout microglia processes

By qualitative observation it was apparent that some processes had little to no mitochondria while other processes had many mitochondria in both mouse and human neocortical microglia (Fig. 3a, b and Supplementary Fig. 3a). To represent this quantitatively the mitochondrial volume in each primary process was plotted against the primary process length, and indeed, primary processes from the same cell had similar lengths but varied in their total mitochondrial volume (Fig. 3c). To determine if the mitochondrial shape differed in these primary processes, we used the mitochondrial volume to separate the primary processes in two groups, designating primary processes as having high (>67.1 μm^3^) or low (<67.1 μm^3^) mitochondrial volume (Fig. 3d). We then compared the shape of the mitochondria in each group. This analysis revealed that the primary processes with low mitochondrial volume had more circular mitochondria compared to processes with high mitochondrial volume that had more elongated mitochondria (Fig. 3e and Supplementary Fig. 3b). Given these data, mitochondria are not uniformly distributed in mouse or human microglia, raising the question of whether there is a functional difference between these processes.

**Figure 3:**
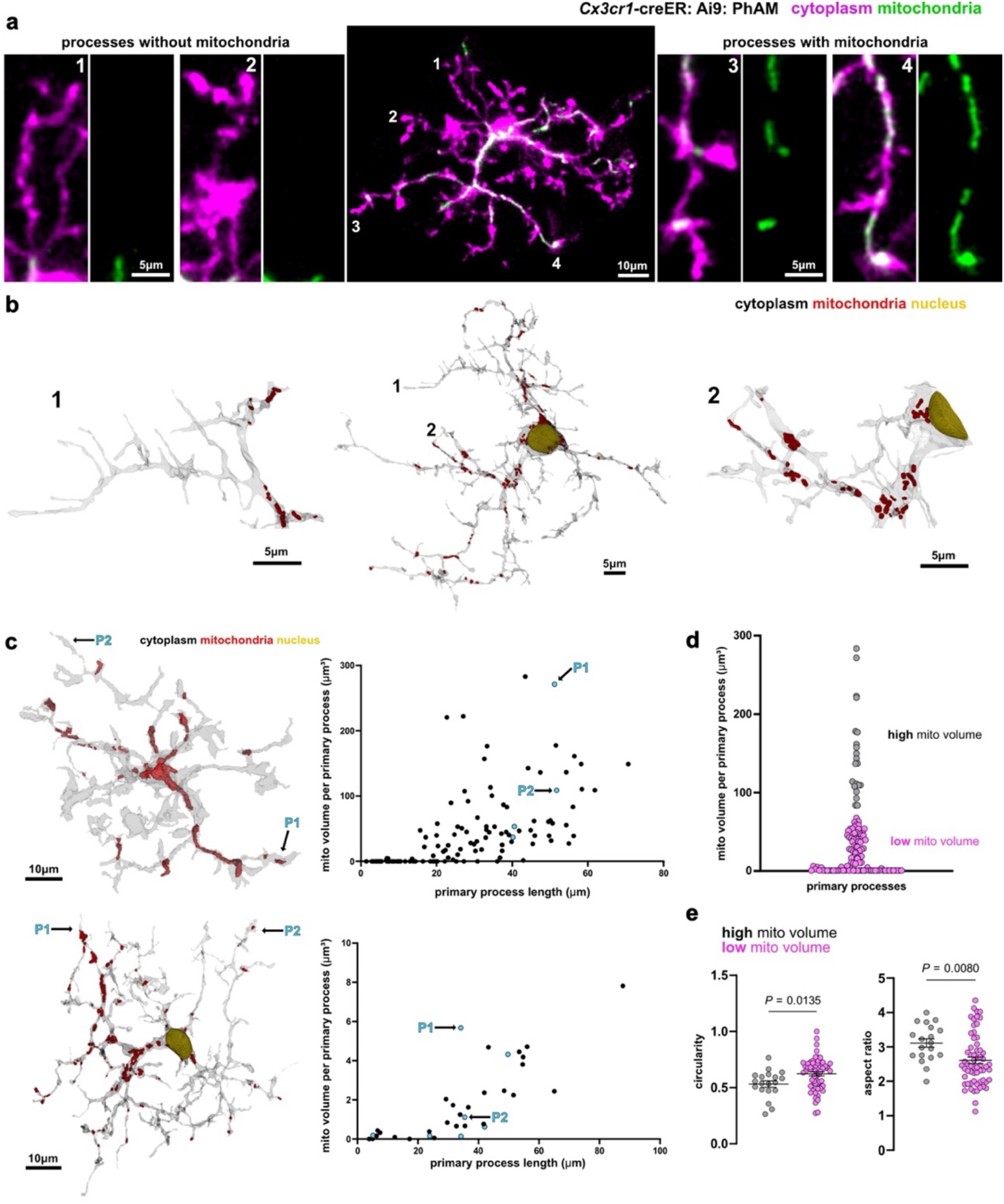
Mitochondria are heterogeneously partitioned throughout microglial processes. **a)** In vivo image of 1 microglia from the mouse cortex (center image) with single regions cropped that represent processes without mitochondria (left images) and with mitochondria (right images) as indicated by number labels 1-4. **b)** 3D rendering of 1 human cortical microglia (center image) from EM segmentations of microglial cytoplasm (gray), mitochondria (red) and nucleus (yellow). Cropped images of processes 1 and 2 represent little to no mitochondria (left image) and many mitochondria (right image), respectively. **c)** 3D renderings of in vivo (top) and EM (bottom) microglia with respective scatter plots of the total mitochondrial volume in each primary process against the primary process length. All dots in the plots represent 1 primary process. Blue dots show the mitochondrial volume in the primary processes of the 3D rendered microglia. Primary processes containing relatively high mitochondrial volume (P1), and low mitochondrial volume (P2) are indicated in the 3D renderings and the graphs. In vivo data, n = 117 primary processes from 14 cells from 6 mice. EM data, n = 34 primary processes from 6 cells from 1 human. **d)** Primary processes from the in vivo data were designated with high (>67.1 μm^3^, 19 primary processes) or low (<67.1 μm^3^, 98 primary processes) mitochondrial volume by k-means clustering (k = 2; n = 117 primary processes from 14 cells from 6 mice). **e)** Mitochondrial circularity and aspect ratio in primary processes with high (n = 19 primary processes) and low (n = 62 primary processes) mitochondrial volume. Primary processes lacking mitochondria were excluded from the analysis (unpaired two-tailed t-test, the line is at the mean ± s.e.m for error bars).

### Microglial branches and processes lacking mitochondria are more motile

To investigate mitochondrial localization and microglial process behavior we next assessed mitochondrial positioning and motility as microglia carry out their canonical surveillance function. To do this, mice were imaged every 1 minute for 30 minutes to enable minute-to-minute tracking of mitochondria in microglia (Fig. 4). First, the stability of 100 branches across 25 cells was assessed. A tracked branch defined as “stable” was present at 0 min and 30 min, and a “lost” branch was present at 0 min but gone at 30 min. Twenty-eight percent of branches were lost and 72% were stable (Fig. 4a). Of all branches assessed, at 0 min, half contained mitochondria (Fig. 4a). Notably, there was a greater percentage of lost branches that did not have mitochondria (Fig. 4a), suggesting that processes lacking mitochondria are more motile. Since much of the surveillant motility occurs at the tips of processes^1,40^ we quantified the number of tips that contain mitochondria and found that only 17% do, supporting the idea that mitochondria are less likely to be present in motile processes (Fig. 4b).

**Figure 4:**
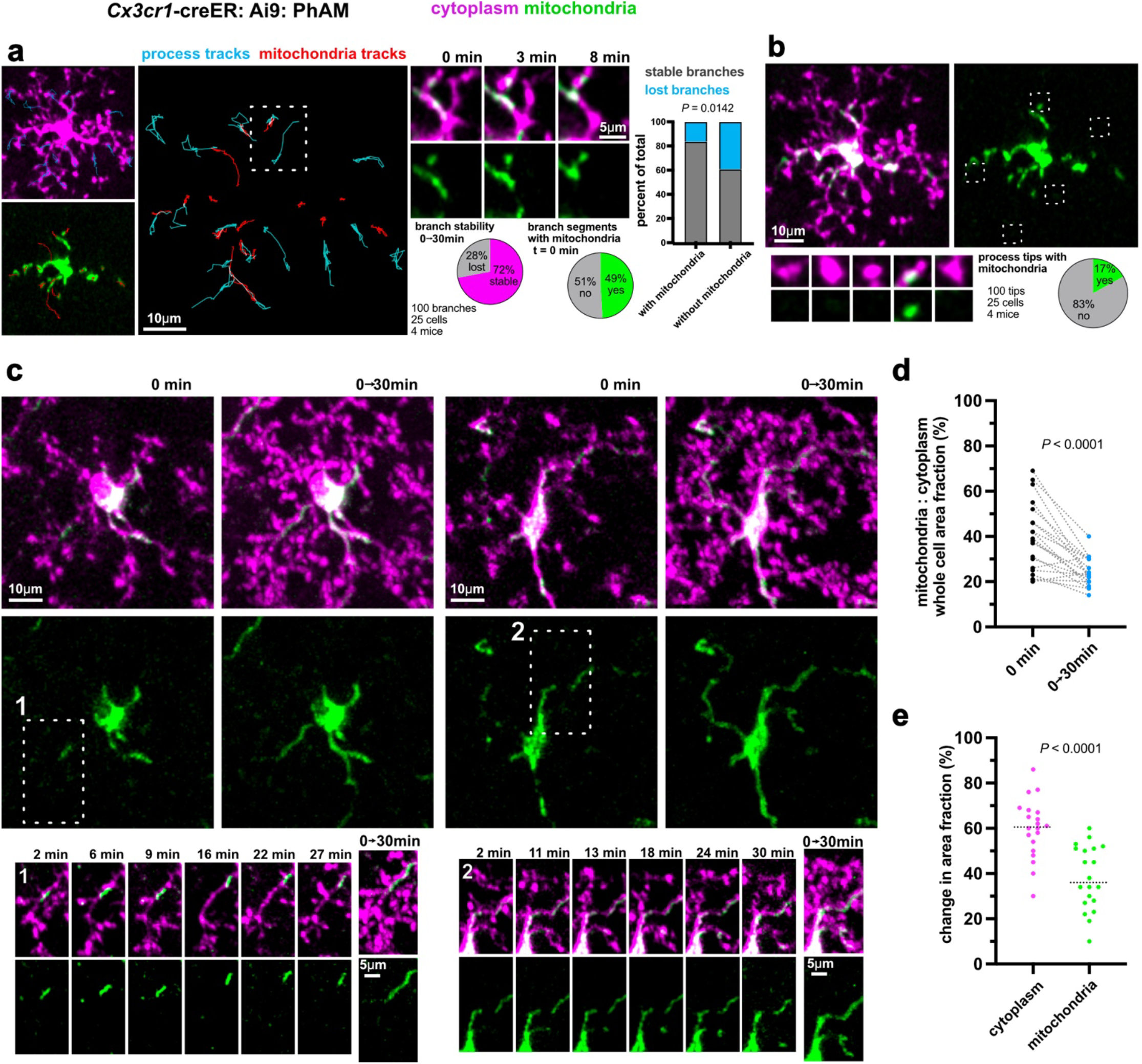
Microglial branches and processes lacking mitochondria are more motile. **a)** Manual tracking of microglia processes (cyan lines) and mitochondria (red lines) over 30 min suggests incongruency between the two. An example of a branch that is maintained and one that is lost with mitochondria present in the branch that is maintained. Pie charts showing the percentage of branches that are stable/lost along with the percentage of branches that contained mitochondria at the beginning of the 30 min time-series. The bar chart compares the stability of the processes that had mitochondria compared to the processes of that did not have mitochondria. (n values indicated in the figure, Fisher’s exact test). **b)** Percentage of microglial process tips that contained mitochondria. **c)** Temporal projections from 0 min to 30 min were used to compare microglia cytoplasmic reorganization relative to mitochondrial motility in an entire cell. The number box insets show examples across the 30 min time-series. **d)** Mitochondria to cytoplasmic area fractions in temporal projections (0→30min) compared to time point 0 min indicating that mitochondria do not move as much as the cytoplasm. **e)** Mitochondria have a lower change in the area occupied relative to the cytoplasm from 0 min to 30 min. For (**d**) and (**e**), n= 25 cells from 4 mice; paired t-tests; dotted lines in (**e**) represent the median.

Next, to determine overall motility we quantified the percent increase in area taken up by the cytoplasm and mitochondria at 0 min and in a temporal projection containing all 30 time points (seen as 0→30min in Fig. 4c). The mitochondria to cytoplasmic fraction significantly decreased in the temporal projection compared to 0 min (Fig. 4c, d). Comparably, the percent change in area fraction of only the cytoplasm (∼60%) is significantly greater than the mitochondrial percent change in area fraction (∼37%) (Fig. 4c, e). Together, mitochondrial motility does not match the microglia process motility across 30 min suggesting that mitochondria are not often associated with processes that are performing tissue surveillance.

### Mitochondrial motility in microglia is not altered by animal anesthesia

Past work has shown that microglia surveillance behavior changes under different physiological states^41,42^, thus we next determined if different physiological states also alter mitochondrial motility in microglia. To do this, we imaged mitochondrial motility in the same microglia as they transitioned from awake to anesthetized conditions. Animals were first imaged awake for 30 min, then switched to isoflurane anesthesia and imaged for another 30 min (Fig. 5a, c). The change in process area fraction showed that microglia were significantly less motile during wakefulness compared to when the mice were under isoflurane anesthesia (Fig. 5a, b, Supplementary Fig. 4), consistent with previous reports^41,42^. However, mitochondrial tracking showed that the mitochondrial displacement and percentage of motile mitochondria were both unchanged in the awake and anesthetized conditions (Fig. 5d and Supplementary Movie 4). Thus, arousal state does not affect mitochondrial motility, again suggesting a disconnect between microglial process surveillance and mitochondrial motility.

**Figure 5:**
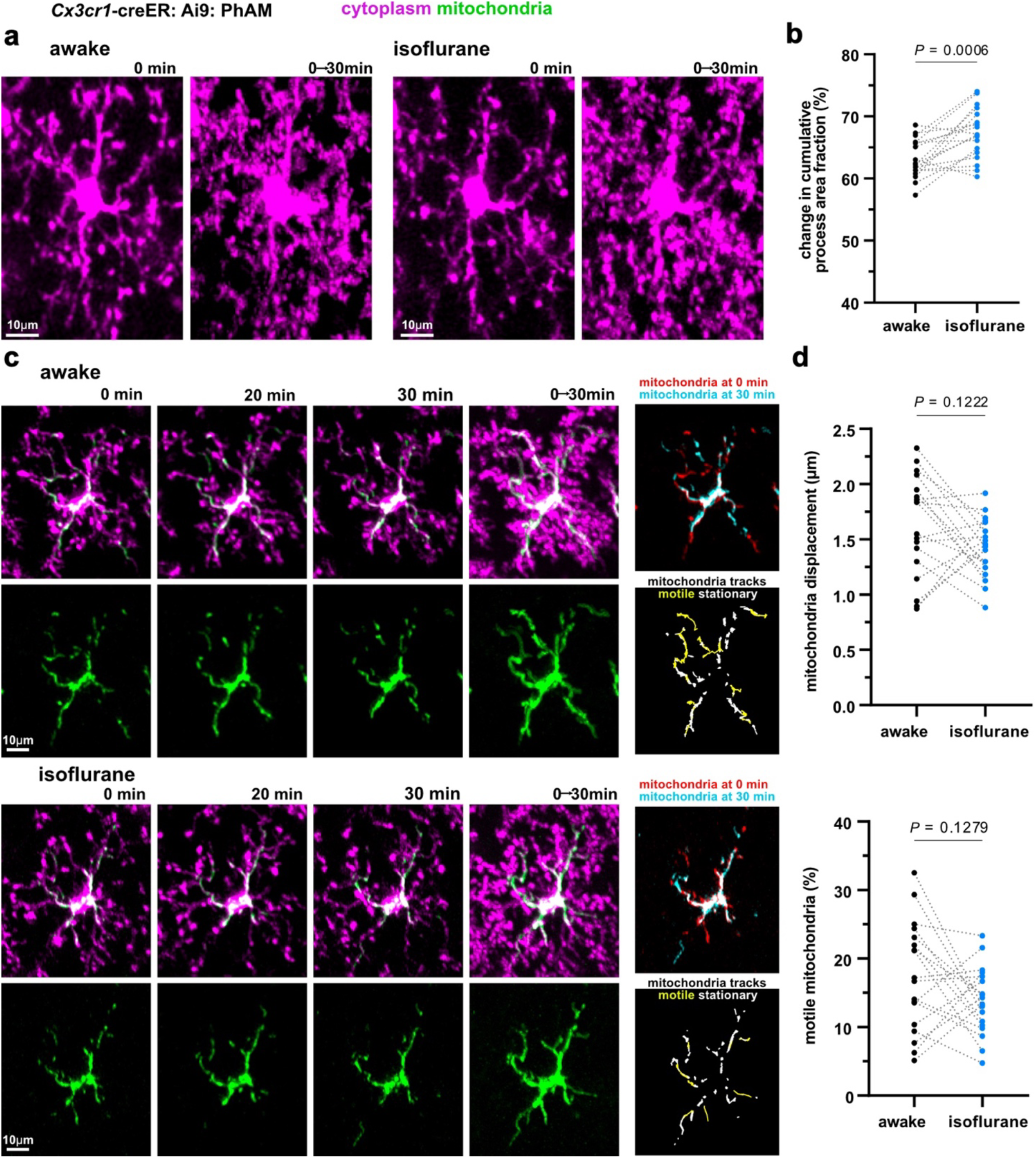
Mitochondrial motility in microglia is not altered by animal anesthesia. **a)** Single time point and temporal projection in vivo images of the same microglial cell when the animal was awake or anesthetized with isoflurane. **b)** The change in cytoplasmic area fraction from the first time point to the 30 min temporal projection shows that microglial process surveillance is greater under the isoflurane condition. **c)** Example time-series showing cytoplasmic and mitochondrial channels in microglial cell imaged in an awake mouse (top) and under isoflurane anesthesia (bottom). Temporal projections show microglial process coverage and mitochondrial coverage over the 30 minutes. The images to the right show the overlap of the mitochondrial channels at 0 min and 30 min and visualization of motile mitochondria (yellow tracks). **d)** Unlike the increase in process motility under isoflurane anesthesia (**b**), mitochondrial displacement (top) and the percentage of mitochondria per cell that move above a displacement threshold of 3 microns (bottom), do not significantly change in the isoflurane condition. For (**a-d**), n = 18 cells from 6 mice; paired, two-tailed t-tests.

### Microglial processes acutely responding to a laser lesion do not contain mitochondria

Given that mitochondria were often not found in surveilling motile microglia processes, we next assessed mitochondrial motility in the context of microglial pathological response to acute tissue damage by a laser lesion. A 30 min time-series was acquired before the laser lesion, and immediately after the lesion a 45 min time-series was taken to capture the microglia and mitochondrial response to the lesion (Fig. 6). To quantify the response, the changes in fluorescence of the cytoplasmic and mitochondrial signals were measured in a region of interest (ROI) around the lesion site, excluding the lesion area (Fig. 6b). As expected^1,3^, multiple microglia processes immediately responded to the laser lesion; these processes arrived around the lesion site within minutes and sustained their presence for hours (Fig. 6c, d, e and Supplementary Movie 5). Unexpectedly, mitochondria were not in the processes that responded to the lesion within the 45 min timeframe (Fig. 6c, d, e and Supplementary Movie 5). To determine if mitochondria arrive later, an image was taken at 1 hour and 6 hours after the lesion. At 1 hour, there was still little to no mitochondria at the lesion site (Supplementary Fig. 4), but at 6 hours mitochondria occupied the processes extending to the lesion site (Fig. 6c, d, e). Therefore, microglia that immediately chemotax to the site of acute tissue damage do not have mitochondria, rather the mitochondria have a delayed arrival into the processes responding to the lesion.

**Figure 6:**
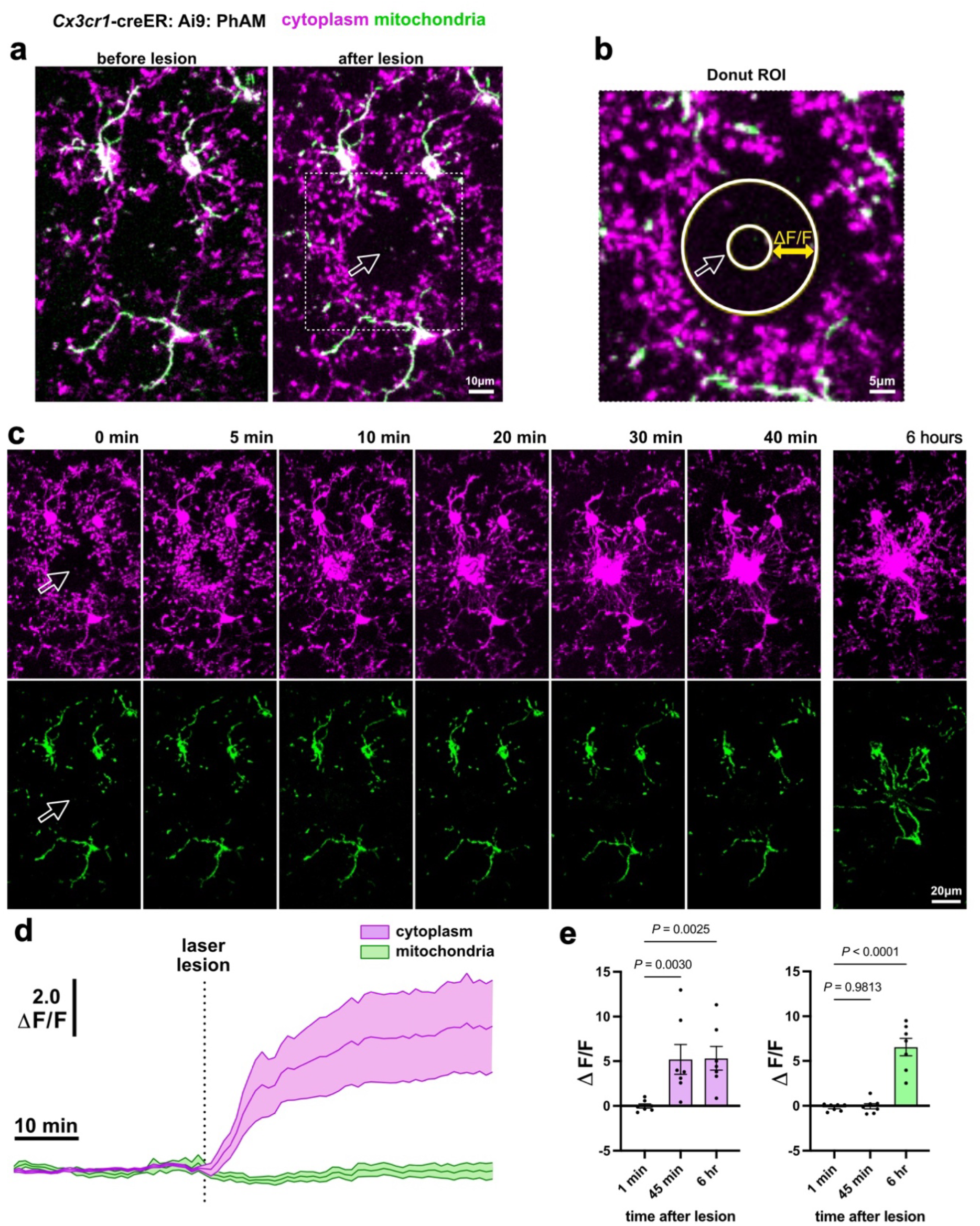
Microglial processes acutely responding to a laser lesion do not contain mitochondria. **a)** A region lacking microglia was chosen for a focal laser lesion (white outlined arrow). After the lesion, a 45min time-series was captured and at 6 hours an image was taken. **b)** A donut ROI was used to measure the change in fluorescence of the microglia and mitochondrial signals before and after the lesion within the defined ROI. **c)** Time-series showing the immediate chemotactic microglia process response to the site of the lesion, but little to no mitochondria response in the same time period. **d)** Fluorescence intensity measurements of the cytoplasmic and mitochondrial channels before and after the laser lesion. **e)** Cytoplasmic fluorescence signals are significantly different from baseline at both 45 min and at 6 hours while mitochondrial signals are not different at 45 min and but show a significant difference at 6 hours after the lesion (for (**d-e**) n = 7 lesions from 6 mice; one-way ANOVA; mean and s.e.m. are represented as error bands in (**d**) or bars in (**e**)).

### Mitochondria are delayed in their arrival to microglial processes engulfing dying neurons

Finally, we investigated the mitochondrial dynamics and subcellular localization when single microglia clear apoptotic neurons. Targeted two-photon chemical apoptotic ablation (2Phatal) involves photobleaching nuclear dye to trigger apoptotic death of neurons (Fig. 7a)^5^. Nuclear condensation can be observed post-2Phatal (Fig. 7a) and the neuronal corpses are cleared by microglia within 24 hours^5,6^. Therefore, 2Phatal serves as a valuable method to track mitochondria as microglia undergo the transition from recognition to early engagement, and then finally engulfment of a targeted neuron. An example of individual microglia actively clearing neuronal corpses 6 hours after 2Phatal can be seen alongside stable microglia that maintained their location and morphology in the parenchyma (Fig. 7a).

**Figure 7:**
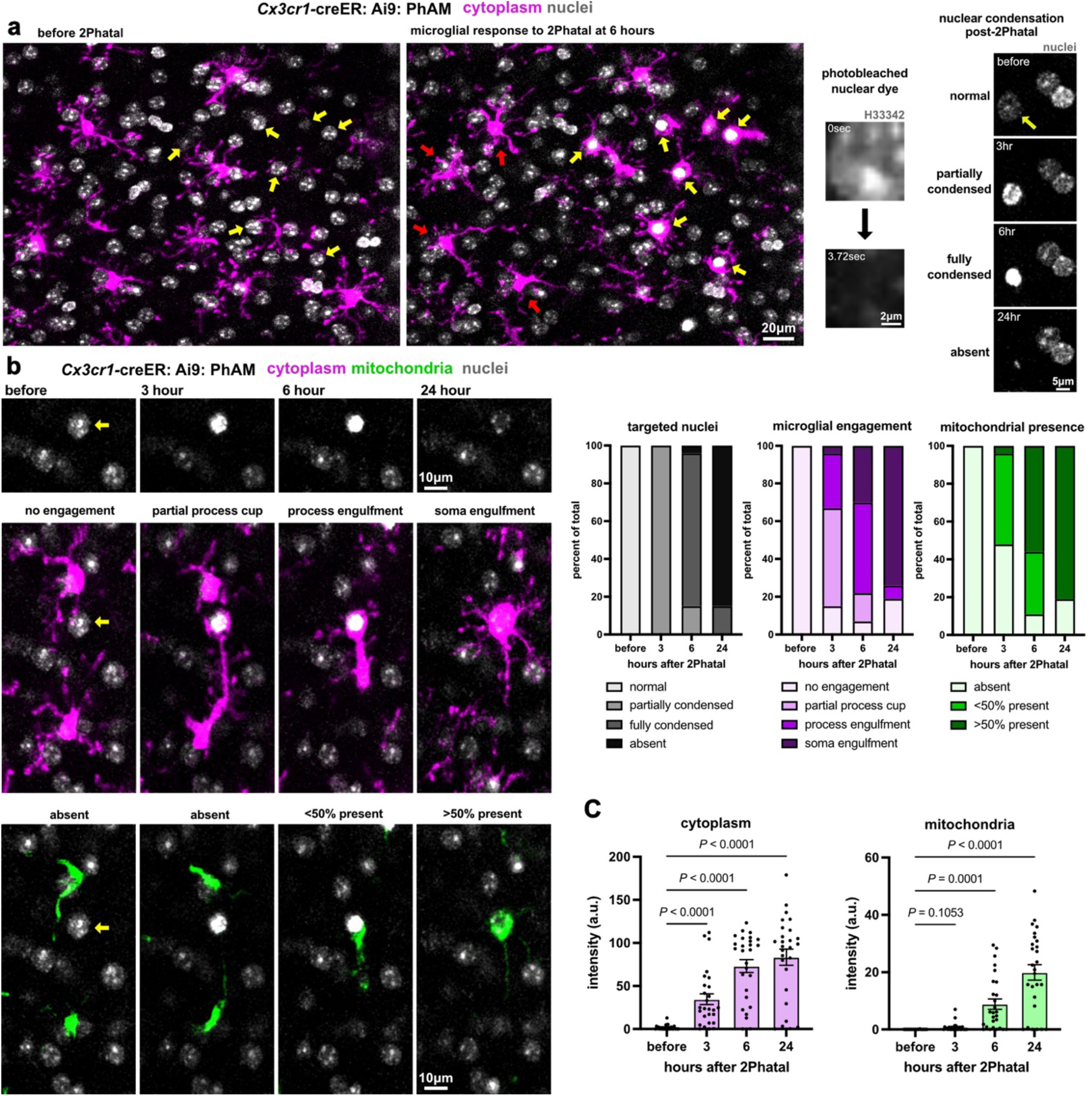
Mitochondria are delayed in their arrival to microglial processes engulfing dying neurons. **a)** Labeling of nuclei (gray) and microglia (magenta) indicating cells that were targeted for 2Phatal (yellow arrows) before and 6 hours after the initiation of cell death. At 6 hours microglia are regularly found to engulf the condensed nuclei of the targeted cells. Nuclei were intentionally targeted on one side of the field of view and microglia that are not engaged in phagocytosis remain ramified (red arrows). The images on the right show an example of nuclear dye photobleaching and the characteristic condensation and clearance of targeted neuronal nuclei. **b)** Time-series showing the engulfment and clearance of a targeted cell (yellow arrow) revealing early cytoplasmic cup formation that lacks mitochondrial labeling followed by full engulfment and microglial repositioning on top of the dying cell. The graphs show scoring of the nuclei (gray), cytoplasm (magenta) and mitochondria (green) at each time point showing that mitochondrial presence lag behind the initial formation of a phagocytic cup. **c)** The fluorescence intensity of the cytoplasm and mitochondrial channels surrounding targeted cells showing that while cytoplasmic signals significantly change at 3 hours and remain elevated, mitochondria show increased abundance at 6 hours. For (**b-c**) n = 27 cells from 6 mice; one-way ANOVA; mean and s.e.m. are represented as error bars in (**c**).

Stages of microglial engagement were tracked at 3, 6, and 24 hours post-2Phatal and defined as 1) no engagement, 2) partial process cup formation, 3) process engulfment, and 4) soma engulfment (Fig. 7b). At 3 hours all nuclei were partially condensed with most microglia forming a partial process cup around the nucleus; of the microglia processes surrounding the nucleus, half were lacking mitochondria, and half had less than 50% mitochondrial presence (Fig. 7b). As the microglia engagement increased to full process or soma engulfment at 6 and 24 hours, there was also an increase in mitochondrial presence within the process or soma engulfing the corpse (Fig. 7b, Supplementary Fig. 6). Fluorescent intensity measurements showed that microglia engagement gradually increased as observed in Figure 7b (Fig. 7c, Supplementary Fig. 6). However, mitochondrial presence in the cytoplasm surrounding the nucleus was not significantly increased until 6 hours after 2Phatal (Fig. 7c). Overall, some mitochondria may arrive 3 hours post-2Phatal, but mitochondrial presence became evident at 6 hours and was sustained when microglia were fully engulfing the dying cell. These findings emphasize a lack of mitochondrial presence in microglia processes during the initial stages of phagocytic engulfment of dying neurons.

## DISCUSSION

In this study, we defined microglia mitochondrial morphometrics and motility in the live intact brain by intravital optical imaging. We found that mitochondria are not uniformly distributed in both mouse and human microglia processes with some processes containing many mitochondria while others had very few. Surprisingly, most of the processes that engage in minute-to-minute surveillance do not have mitochondria. Likewise, as microglia respond to a laser lesion, the processes that immediately chemotax to the lesion site do not have mitochondria, nor do the processes that initiate the initial stages of engulfment of an apoptotic neuron. Rather, the mitochondria have a delayed arrival into the microglia processes responding to injury. Additionally, mitochondrial motility was unaltered by the arousal state (awake or anesthetized) of the animal even though the microglial process motility was affected. Together, this study presents the heterogeneity of mitochondrial subcellular distribution in microglia during surveillance, chemotaxis, and phagocytic responses.

A broad look at mitochondrial distribution in neocortical microglia showed that more mitochondria resided at the cell center and decreased towards the tips of the processes consistent with previous findings from the murine retina^32^. These observations contradict previous findings from in vitro studies as cultured microglia generally have more fragmented mitochondria spread evenly throughout the cell^28–30^. Our analysis also found that human cortical microglia showed similar mitochondrial partitioning and morphometrics, namely that some microglia processes are completely devoid of mitochondria (Fig. 1 and Fig. 2). Together, these data emphasize the importance of defining mitochondrial dynamics while microglia are maintained in their native microenvironment, as culture studies do not closely model the complex CNS environment.

Mitochondria have phenotypic and proteomic differences across organs^11^. However, the uneven distribution of mitochondria in microglia with some processes containing little to no mitochondria and others being filled with mitochondria (Fig. 3) contrasts the mitochondrial distribution seen in other neural cells in the brain. For instance, mitochondria occupy almost all the complex arborizations of neurons^43,44^ and astrocytes^45^. As neurons consume most energy in the body and astrocytes are highly involved in their metabolic fitness it makes sense that mitochondria would be situated in all parts of the cell. We expected the radical morphological changes microglia undergo during surveillance, chemotaxis, and phagocytosis would require mitochondrial support for the energetic demand of those processes. However, we found that more motile microglial processes performing surveillance were less likely to contain mitochondria compared to the stable processes (Fig. 4). Another brain cell that undergoes extensive morphological changes is the oligodendrocyte precursor cell (OPC) as it differentiates into a myelinating oligodendrocyte^46^. At the stage when OPCs begin to form myelin, the arborizations are filled with mitochondria, but when the oligodendrocyte matures, mitochondria are concentrated in the soma and are mostly lost in the myelin sheaths^47^. The OPC to oligodendrocyte differentiation process takes days to weeks while microglial morphological changes can occur in a matter of minutes to hours. Therefore, the reorganization of internal cellular architecture may require time to manifest as microglial morphology changes. Indeed, although microglial processes reached the site of acute tissue damage (within 10 min, Fig. 6) or to an apoptotic neuron (within 3 hours, Fig. 7), the mitochondria did not arrive within those processes until several hours after the injury (Fig. 6 and Fig. 7). Other studies show that mitochondria follow or occupy tips of regenerating axons^48^, the leading edge of migrating cells^49–51^, and in the tips of filopodia^52^. Whereas we rarely see mitochondria in the tips of surveillant processes or in the microglial processes that are performing chemotaxis to the site of injury (Fig. 4 and Fig. 6). Thus, microglia orchestrate mitochondrial subcellular distribution and motility differently from other migratory cells or other cells undergoing shape changes.

One explanation for the differences between cell types is that actin polymerization is mainly thought to drive microglial surveillance and the immediate chemotaxis to the site of acute tissue damage^4,53^. However, mitochondria are primarily transported along microtubules^13^. Given that, it is likely that the timing of mitochondrial arrival into the processes during the final stages of full engulfment is linked to actin vs microtubule cytoskeletal rearrangements occurring during the microglial injury response. In the developing zebrafish, centrosome orientation and microtubule organization determine which microglia process will engulf an apoptotic neuron^54^. These experiments revealed that it takes time for the centrosome to travel through the cell and enter the branch that will perform phagocytosis^54^. This supports the hypothesis that there is a selectivity to internal structure reorganization and functional heterogeneity in the microglia processes of a single cell. Future studies can explore the interplay between mitochondrial dynamics and cytoskeletal organization as microglia carry out their multifaceted functions.

It is important to acknowledge that by optically imaging mitochondria we are limited by resolution capabilities. Therefore, the shape of mitochondrial networks rather than individual mitochondria were analyzed. By aspect ratio, we found that mitochondria in the processes were more elongated compared to soma mitochondria, but the circularity measurement showed no difference (Fig. 1). The shape difference seen by aspect ratio was likely undetected by the circularity measurement since different features of the object are considered by these two methods to calculate shape (see *Mitochondrial morphometric analysis* in methods). The ultrastructure resolution from the electron microscopy (EM) 3D reconstructions provided corroborating evidence that mitochondria are more abundant but smaller in the processes, while in the soma, there are fewer mitochondria that are larger (Fig. 1 and Fig. 2). EM allows for high resolution spatial reconstruction, but the tissue fixation likely affects microglial and mitochondrial morphologies^55^, explaining why many of the mitochondria in the EM dataset were more fragmented compared to the mitochondria imaged in vivo.

Innate immune cells undergo a metabolic shift from oxidative phosphorylation to glycolysis in response to injury to effectively host a response^56^. Past work in vitro has shown that microglia switch metabolic profiles when exposed to inflammatory stimuli^28–30,57^. A challenge of studying microglia in the intact brain is the inability to thoroughly investigate microglial metabolism alongside mitochondrial dynamics and how these change during surveillance, chemotaxis, and phagocytosis. One study partially circumvented this limitation by removing a metabolic protein needed for the first step of glycolysis^58^, hexokinase2 (HK2), specifically in microglia^33^. They found that microglial process motility, velocity, and response to a focal lesion were all slowed in the live mouse with HK2 knocked out^33^. These in vivo findings were complemented with metabolic profiling in primary cultured microglia, revealing a drop in both glycolysis and oxidative phosphorylation in the microglia with HK2 knocked out, indicating that HK2 loss left microglia energetically depleted^33^. There has been growing evidence that metabolic dysregulation and mitochondrial dysfunction lead to impaired phagocytic capacity of microglia contributing to neurodegenerative disease pathology and developmental abnormalities^31,34–36,59,60^. Future studies examining microglial bioenergetics in development, neuroinflammation, and neurodegeneration would benefit from also investigating mitochondrial morphometrics and dynamics using the approaches developed and employed here.

## Acknowledgments

We would like to thank past and current members of the Hill and Hoppa laboratories at Dartmouth College for their valuable feedback and support throughout the project. We also thank Andrew D. McCray for helping with data analysis. This study was supported by National Institutes of Health grant R01NS122800, diversity supplement R01NS122800-S1, and the Esther A. & Joseph Klingenstein Fund and Simons Foundation to R.A.H; and the American Heart Association grant, 23PRE1018862, https://doi.org/10.58275/AHA.23PRE1018862.pc.gr.161154 to Xh.B.

## Author contributions

A.N.P and R.A.H conceived, designed and performed all the experiments, and most of the data analysis, quantification, and writing of the manuscript. Xh.B performed the analysis and quantification for the microglial process and mitochondrial motility, and contributed to data representation and writing pertaining to these analyses. M.E.D contributed to the analysis and quantification of microglia and their mitochondria from the publicly available H01 electron microscopy dataset. A.N.P and R.A.H secured funding and R.A.H supervised the study.

## METHODS

### Animals

The triple transgenic mouse model used in this study, *Cx3cr1*-creER: Ai9: PhAM, was generated by breeding the following mouse lines together: *Cx3cr1*-creER (JAX strain# 020940)^61^, floxed tdTomato (JAX strain# 007909)^62^, and PhAM (JAX #018385)^63^. This line has conditional expression of cytoplasmic localized tdTomato (Ai9) and mitochondrial matrix localized Dendra2 (PhAM) to all *Cx3cr1+* cells, which are predominantly microglia in the central nervous system. Animals used for this study were heterozygous for creER to prevent knock out of *Cx3cr1*. To mediate cre recombination, mice were administered an intraperitoneal (i.p) injection (0.05 mL) of tamoxifen (Sigma-Aldrich #T2859-1G) dissolved in corn oil (20 mg/mL) every other day for 5 days (total of 3 injections) at weaning. Both male and female adult mice were used at 3-6 months for the experiments^64^. All animal procedures were approved by the Institutional Animal Care and Use Committee at Dartmouth College (protocol number: 00002158). All animals were housed in a temperature and humidity-controlled vivarium in a 12 hr light/dark cycle with ad libitum access to food and water.

### Surgical procedures

Cranial window surgeries were performed over the somatosensory cortex as previously described^47,65,66^. Briefly, animals were anesthetized with an i.p. injection of ketamine (100 mg/kg) and xylazine (10 mg/kg) solution. Carprofen analgesic (50 mg/kg) was administered subcutaneously before, after, 24 hr after, and 48 hr after the surgery. A custom-made head plate was glued to the exposed skull and then a craniotomy was performed to remove a circular portion of the skull (3 mm × 3 mm). For 2Phatal experiments, Hoechst 33342 nuclear dye (1:175) was directly applied to the exposed brain. The brain was sealed with a no. 0 cover glass allowing for intravital imaging. Experiments were performed 3-5 weeks post-surgery to allow the surgery to heal and for inflammation to subside. However, Hoechst 33342 optimally labels nuclei 24 hr after topical application to the brain, therefore, we performed the 2Phatal experiments 24 hr after the cranial window surgery.

### Intravital imaging

While the mouse was still anesthetized from the cranial window surgery, high resolution images were acquired on an upright confocal microscope (Leica SP8) with a 20X water immersion objective (Leica NA 1.0). All other imaging was performed on an upright laser-scanning 2-photon microscope (Bruker Ultima) with a 20X water immersive objective (Zeiss NA 1.0). The cytoplasmic tdTomato was excited by 552 nm and 1040 nm laser wavelength on the confocal and 2-photon, respectively. The mitochondrial Dendra2 was excited by a 448 nm and 920 nm laser wavelength on the respective microscopes. The Hoechst 33342 nuclear dye was excited by a 775 nm laser wavelength on the 2-photon. All Z-stacks were taken over a depth of 50-60 μm with a 1.5 μm step size between stacks. The imaging depth spanned layers 1 and 2 of the somatosensory cortex. All time-series were acquired by taking a Z-stack image every 1 min for a total of 30, 45, or 60 min. For anesthetized imaging sessions isoflurane anesthesia mixed with room oxygen was delivered to the animal via the SomnoSuite, Low-Flow Anesthesia System. Initially, 3.5% isoflurane was used to induce anesthesia, then anesthesia was maintained at 1.5% isoflurane while imaging. Awake imaging experiments were performed by head fixing the mouse by the head plate in the Neurotar’s Mobile HomeCage that allows the mouse to move ad libitum on an air table.

### 3-dimensional segmentation and rendering from optical images

High resolution confocal images were used to segment microglial cytoplasm and their mitochondria. The image size was 2048 × 2048 pixels (pixel size 180 nm × 180 nm), and 1-3 positions were acquired per animal. A total of 14 microglia from 9 mice were segmented. The parameter to choose the microglia to segment was that all cell compartments had to be within the Z-depth imaged. Aside from that, the cells to segment were randomly selected (1-3 cells per animal).

The Volume Annotation and Segmentation Tool (VAST Lite, version 1.4.1)^67^ was used to segment the cytoplasmic (tdTomato) and mitochondrial (Dendra2) signal in 3D. Images were preprocessed in FIJI^68^: the tdTomato and Dendra2 channels were split, smoothed, the look up table (LUT) changed to Grays, and converted from an 8-bit to RGB Color. The images were then imported to VAST to complete the segmentations. For each cell, first the cytoplasm was segmented to define the soma, processes, and primary process compartments. Then the mitochondrial networks were segmented and designated to the respective compartment they resided in. Lastly, with VAST we exported the following from the completed segmentations: 1) the cytoplasmic and mitochondrial volumes and 2) uncompressed TIFF images at mip level 0 (indicates the voxel resolution in X, Y, Z: 180 nm, 180 nm, 1500 nm). Both were later used in mitochondrial morphometric quantifications done in FIJI or Excel. The raw values for mitochondrial volume from the optical images were likely overestimated due to limits in axial resolution with optical fluorescence microscopy in addition to the inability to always resolve and separate single mitochondria from mitochondrial networks as mentioned in the discussion. Therefore, direct comparisons of mitochondrial volume between the optical and EM datasets cannot be made. This does not impact other comparisons as ratio-metric or within cell comparisons are accurate.

To 3D render the microglial cytoplasm and mitochondria, MATLAB (version 24.1.0)^69^ and VAST were used to export OBJ files of the cytoplasmic and mitochondrial segments at mip level 0. The OBJ files were used in 3D Slicer^70–73^ to create the 3D renderings (see examples in Fig. 1e and Supplementary Fig, 1c).

### Human electron microscopy dataset analysis

To investigate mitochondria in microglia across species we used a publicly available human electron microscopy (EM) dataset (H01) available online at http://h01-release.storage.googleapis.com/landing.html^38^. This dataset provides a complete cellular and vascular reconstruction of 1 cubic millimeter of the temporal cortex, the largest reconstruction of the human brain to date^38^. The brain tissue was donated from a 45-year-old female who underwent an epilepsy-related resection surgery^38^. To access and remove the damaged tissue, surgeons needed to also remove a healthy, non-pathological section, a portion of which constitutes the H01 dataset^38^.

The tissue was prepared as described^38^, sectioned at ∼30 nm thickness, and imaged using multibeam scanning electron microscopy with a resolution of 4 × 4 nm^2^. This group identified the cell types and segmented their shape in 3D, which can be viewed in Neuroglancer^38^. We distinguished microglia from OPCs by their nuclear ultrastructure and cell cytoplasmic morphology^39^. Using Neuroglancer, we chose 6 microglia from cortical layer 5 to further segment their mitochondria. We used VAST^67^ to open the H01 dataset, import the coordinates of our cells of interest taken from Neuroglancer, and fill the microglia cytoplasm that had already been automatically segmented^38^. We further segmented other microglial compartments including the soma, processes, primary processes, nucleus, and mitochondria. Mitochondria were designated to their respective compartment. Similar to the fluorescent segments, we used VAST to export the following from the specific bounding area of the EM segmentations: 1) the cytoplasmic, nuclear, and mitochondrial volumes and 2) uncompressed TIFF images at mip level 4 (X, Y, Z: 128 nm, 128 nm, 132 nm). A higher mip level lowered the resolution such that images could be exported in a timely manner. These exports were later used for mitochondrial morphometric quantifications done in FIJI or Excel.

### Mitochondrial morphometric analysis

Morphometrics in this study encompass mitochondrial content, shape, and subcellular distribution. Starting with content, all volume exports from the 3D segmentations were used to determine the mitochondrial to cytoplasmic volume fraction (%) and the total or average mitochondrial volume (μm^3^) in the whole microglia and/or by microglial compartment (soma and processes). Additionally, mitochondrial network number and the number of individual mitochondria were manually counted while segmenting the optical (mouse) and EM (human) datasets, respectively. For the optical dataset, if a mitochondrial network spanned more than one compartment, its’ volume was split to represent the compartment it resided in, but it was counted as 1 network. For the EM dataset, if a mitochondrion spanned more than one compartment, it was assigned to the one it most occupied.

From the optical segmentations, mitochondrial shape was determined by circularity and aspect ratio: two different calculations to determine how circular or elongated an object is. A value equal to 1.0, from both circularity (4pi*(area/perimeter^2^)) and aspect ratio (width/length), represents an object that is a perfect circle. Therefore, as the value moves further from 1.0, the object becomes more elongated. To quantify the mitochondrial shape, mitochondrial segments from each microglia were opened in FIJI, a maximum intensity projection was made, default threshold applied, then the “Analyze Particles” command was used to automatically calculate circularity and aspect ratio from the shape descriptors. Circularity and aspect ratio consider different features of the object. Using both methods ensures a thorough analysis, as one calculation might not detect a shape difference, while the other does.

Multiple analyses were used to dissect the subcellular distribution of mitochondria in microglia. First, a sholl analysis was done on the cytoplasmic and mitochondrial segments (from optical and EM datasets) separately. Sholl is used on cells with a branched morphology to determine the complexity of the branching. It does this by counting the number of intersections that branches make on evenly spaced concentric circles that radiated out from the cell center. With sholl, we wanted to determine how mitochondria radiate from the cell center relative to the microglial branching complexity. To perform sholl, maximum intensity projections were made from the cytoplasmic and mitochondrial segments in FIJI, a default threshold was applied, a straight line was drawn from the cell center to the edge of the image, and the “Neuroanatomy” plugin was used to execute sholl. Sholl parameters for the optical dataset were set to 1 μm for the starting radius and step size with a 70 μm ending radius. Given the increased spatial resolution of the EM dataset, we used the following sholl parameters: 0.5 μm for the starting radius and step size with an 80 μm ending radius. Results were graphed as number of intersections against the distance from the cell center. Of note, to exclude contributions from the soma, radii 1 – 8 μm were excluded from the sholl graphs (Fig. 1 and 2) since the average soma radius of microglia is around 8 μm in the optical and EM datasets (data not shown).

The second set of analyses determined the subcellular distribution of mitochondria along the length of individual primary processes in 10% bins. To measure the primary process lengths, the optical 3D segments (from the cell soma and the primary process cytoplasm and mitochondria) were first maximum intensity projected and merged into one image in FIJI. In the merged image, only the cytoplasm and soma channels were made visible. The mitochondrial channel was off to ensure the researcher was blind to the mitochondria location along the primary branch length. Next, a 5-pixel segmented line tool was used to trace the longest branch in the primary process from soma end to tip end such that the shortest possible path was traced. Then, the “Plot Profile” command was executed in reference to the mitochondrial channel which provided the primary process length (μm) and information (from gray value) on where mitochondria specifically reside along that length. To interpret this information, the length of all the primary processes were normalized to each other by considering the length as 0-100% (soma = 0% and tip of process = 100%) and quantifying the mitochondrial occupancy in 10% bins along the length of the primary process.

The third set of analyses further described mitochondrial subcellular distribution in primary processes by scatter plotting the mitochondrial volume in each primary process (μm^3^) by the primary process length (μm) with both the optical and EM datasets. The primary process lengths of the human EM microglia were measured with plot profile as described above. Primary process length, not cytoplasmic volume, was used in these analyses to distinguish long processes from short processes. Cytoplasmic volume was misleading because some short processes had greater volume than longer and thinner processes.

Lastly, Excel was used to perform a k-means cluster (k = 2) using the mitochondrial volume per primary process. This computationally defined two groups that contained either high mitochondrial volume (>67.1 μm^3^) or low mitochondrial volume (<67.1 μm^3^). Within those 2 groups we determined the circularity and aspect ratio of the individual mitochondrial networks in each primary process (excluding the primary processes lacking mitochondria). The goal of this analysis was to determine if mitochondrial shape in a microglia process differs depending on the volume of mitochondria occupying that process. We used the optical mouse dataset for this analysis given the abundance of primary processes analyzed.

### Mitochondrial presence during microglial surveillance

To determine mitochondrial presence during microglial surveillance, animals were imaged under isoflurane anesthesia 3-5 weeks after the cranial window surgery (see *Surgical Procedures* and *Intravital Imaging*). A 30 min time-series was captured from 4 mice. In FIJI, the time-series was 3D Drift Corrected based on the tdTomato channel, then 25 microglia were selected also using the tdTomato channel. Each microglial cell was cropped, smoothed, then maximum intensity projected from the 5 slices above and 5 slices below the center of the microglial soma. Then branches (3-6 per cell) and process tips (3-6 per cell) were selected (using the tdTomato channel) from each microglia and the following analyses were performed.

We defined the branches as “stable”, present at 0 min and 30 min, or “lost”, branch was present at 0 min but gone at 30 min. Stable branches were considered less motile than the lost branches. With this we determined the percentage of branches that were stable or lost. To confirm that the branches selected had no difference in mitochondrial occupancy at timepoint 0 min, we determined the percentage of branches that contained and did not contain mitochondria. Then, we determined the percentage of branches containing mitochondria in the stable branches vs. the lost branches. Given that much motility occurs at the tips of the processes we overlayed a 5 × 5 μm^2^ region of interest (ROI) around the tip’s center and determined the percentage of tips that contained or did not contain mitochondria.

Next, we quantified the area covered by the cytoplasmic and mitochondrial signal across the 30 min time-series by temporally projecting all the 30 images together (designated as 0→30 min). We visualized process and mitochondrial motility from manual tracks made in the FIJI’s TrackMate plugin. An increase in area fraction indicates motility, therefore, we aimed to quantify the motility by these methods to compare microglial process and mitochondrial motility during surveillance. Each cropped cell was further processed in FIJI: the cytoplasmic and mitochondrial channels were split, bleach corrected, “Li” thresholded to make a binary image with minimal background signal, and finally temporally projected. From this, area fraction from the cytoplasmic and mitochondrial channels was determined from the 0 min image and the temporal projection 0→30 min. We determined the percent of mitochondrial to cytoplasmic area fraction, and the percent change in area fraction for each signal separately.

### Awake to isoflurane anesthetized imaging

Awake to anesthetized imaging was performed 3-5 weeks after the cranial window surgery (see *Surgical Procedures* and *Intravital Imaging*). First, the animal was acclimated to the Neurotar Mobile HomeCage for 30 min, then a 30 min time-series was captured. Next, the same animal was removed from the awake imaging set-up and anesthetized. We allowed the mouse to acclimate to the anesthesia for 30 min before locating the same position that was imaged during wakefulness. Then, a 30 min time-series was captured. Therefore, two 30 min time-series were acquired from the same animal in the same position: first during wakefulness and then during anesthesia allowing us to compare the same microglia and their mitochondria under different arousal states.

### Motility of microglial processes and mitochondria

For analyzing the mitochondria and processes’ dynamics in awake and isoflurane anesthetized conditions, first, the Z-stacks were registered to correct for the Z-drift with reference to the cytoplasmic channel. The cells to be analyzed were randomly selected with reference only to the first time point and the analyzer was blinded to the experimental condition. The cells were cropped, smoothed, and used to create Z-projections which were later thresholded similarly for the mitochondria and cytoplasmic channel. To measure the processes’ dynamics, Z-projections of the first and last time points as well as of all the time points along the 30 min time-series were created for the cytoplasmic channel. Area fraction in the cropped region was measured for the first time point as well as for the temporal projections and the change in area fraction between the two was calculated.

Mitochondria motility analysis was done only for the mitochondria in the processes of the selected cells and mitochondria present in the soma or other cells in the environment were cleared out. The thresholded images were then run in TrackMate^74^ with the following settings: the LoG detector was selected, and the estimated object diameter was set at 3, quality threshold at 1, and sub-pixel localization was checked; the simple LAP tracker was selected and the linking and gap-closing max distance was set at 3 microns, and gap-closing max frame gap at 2. The tracks with less than 5 spots were excluded and displacement was collected for each of the automatically tracked mitochondria networks. To correct for any remaining drift caused by imaging and sample motion artifacts, first, the mitochondria track that traveled the smallest distance along the 30 min time-series was identified. It was assumed that any displacement collected from this track was due to drift and therefore all the other displacement values were corrected accordingly based on this track. All the above steps were kept consistent for both awake and isoflurane conditions. Lastly, all the mitochondria displacement values of all the tracks analyzed in both conditions were combined to calculate the mean and standard deviation. Their sum (rounded up to 3 microns) was determined to be the displacement threshold above which the mitochondria tracks were considered motile.

### Laser lesion imaging and analysis

The laser lesion experiment was performed in awake animals using the Neurotar Mobile HomeCage 3-5 weeks after the cranial window surgery (see *Surgical Procedures* and *Intravital Imaging*). The laser power was increased to ∼75mW for the lesion. The high laser power used for the lesion causes cell rupture and spillage of ATP into the parenchyma^3^. ATP then triggers a chemotactic response from microglia via the transmembrane purinergic P2Y12 receptor^75^. This method of applying focal tissue damage serves as a model of acute inflammatory response by microglia. Before the lesion was induced a 30 min time-series was acquired to capture baseline surveillance. In the same position, a laser lesion was induced in an 8 × 8-pixel ROI in an area with little to no fluorescent signal, then immediately after the lesion a 45 min time-series was captured. This allowed us to observe the microglial transition from baseline surveillance to rapid chemotactic response to the lesion and assess the mitochondrial positioning and motility throughout this process. Additionally, the same position was imaged at 1 and 6 hours after the lesion to observe the progression of mitochondrial dynamics as microglia work to clear the debris. The same position was identified at each timepoint using the vascular morphology and cellular patterning.

To determine when microglial processes and mitochondria arrive to the laser lesion, the change in fluorescence over time was quantified in the region surrounding the lesioned area. To do this the 30 min and 45 min time-series were preprocessed as follows: a maximum intensity projection was made, the tdTomato and Dendra2 channels were each bleach corrected, then the projection was corrected for 3D drift referring to the Dendra2 channel, the tdTomato and Dendra2 channels were split, then the lesion location was identified. Next, autofluorescence produced from the lesion was removed by clearing the all tdTomato and Dendra2 signal in a 16 × 16-pixel ROI (double the size of the original 8 × 8-pixel lesion), then the “Li” threshold was applied to the projections to create a binary image stack of each channel and remove background signal. Next, a 50 × 50-pixel ROI was centered around the cleared area, creating a donut ROI (Fig. 6b), and the “Time Series Analyzer V3” FIJI plugin was used to quantify the average fluorescent intensity around in the donut ROI over time. The average fluorescent intensities across the 30 min times-series were averaged for each position and channel (tdTomato and Dendra2) imaged. This average served as the baseline fluorescence for the position, and it was used to calculate the change in fluorescent intensity minute by minute for the 30 min before and 45 min after the lesion was induced. The change in fluorescent intensity was also quantified from a single 6 hr time-point to determine the possible progression of mitochondrial arrival to the lesion site.

### 2Phatal imaging and analysis

Targeted two-photon chemical apoptotic ablation (2Phatal) experiments were performed in awake animals using the Neurotar Mobile HomeCage 24 hours after the cranial window surgery (see *Surgical Procedures* and *Intravital Imaging*). 2Phatal enabled us to trigger apoptotic death of single neurons^5^ in layer 2 of the somatosensory cortex. We observed the microglia mitochondrial response to this non-inflammatory injury^7–9^ as a single microglia recognized the apoptotic neuron and initiated phagocytosis of the neuronal corpse^5,6^. Neuronal nuclei were identified by their granular morphology and large size^5^. 2Phatal was induced as previously described^5,65^. Briefly, an 8 × 8-pixel ROI was placed over the Hoechst 33342 labeled neuronal nucleus, the dwell time was increased to 100 μs, the laser intensity was increased, and then the nuclear dye was laser ablated for 3.2 s. This damages the DNA and triggers apoptosis^5^. One position per animal was captured and within each position, 1-8 neuronal nuclei were targeted. Images were taken before, 3 hr, 6 hr, and 24 hr after 2Phatal. The same position was identified at each time point using the vascular morphology and nuclear patterning. This is a valuable non-inflammatory cell death method that will enable visualization of how the mitochondrial dynamics interplay with the microglial process dynamics when clearing a single neuron.

First, we qualitatively assessed the nuclear condensation, microglial engagement, and mitochondrial presence at each timepoint imaged (before, 3 hr, 6 hr, and 24 hr). Nuclear condensation was defined as either normal, partially condensed, fully condensed, or absent. Microglial engagement was defined as no engagement, partial process cup, process engulfment, or soma engulfment. Mitochondrial presence was defined as absent, <50% present, or >50% present. These scorings of engagements were individually graphed to show the percentage of total cells presenting the specific nuclear, cytoplasmic, and mitochondrial phenotypes before and after 2Phatal.

For a quantitative assessment of the microglial cytoplasmic and mitochondrial response to clearing an apoptotic neuron, we determined the average fluorescent intensity at each time point (before, 3 hr, 6 hr, and 24 hr) imaged post-2Phatal. To perform this analysis in FIJI, a 20 × 20 μm^2^ circular ROI was centered the 2Phatal targeted nuclei to produce a cropped image for each nucleus at each time point. From the cropped images, the channels (tdTomato and Dendra2) were split, and a default threshold was applied to each image. Then, a 20 × 20 μm^2^ circular ROI was overlayed on each image and the mean gray value was effectively quantified around the immediate vicinity of the target nucleus across all timepoints and in both the cytoplasmic and mitochondrial channels.

### Statistical analysis

All statistical analyses were performed using GraphPad Prism (version 10.3.0) or Excel. No statistical methods were used to predetermine sample size, and all data was assumed to have a normal distribution. Sample sizes were based on those reported in previous publications. Statistical tests, cell numbers, and animal numbers are indicated in the figure legends for each experiment. The parameters to select the cells analyzed in this study were based on the quality of the images and that all cell compartments were within the imaged Z depth. The cells for the motility analyses were randomly selected with reference to the first time point of the time-series, and the data was blinded for awake and isoflurane anesthetized conditions. The cells analyzed from the EM dataset were selected based on the microglial identification criteria. Throughout the study, the analyzer only referred to the cytoplasmic channel, and not the mitochondrial channel, when defining ROIs to prevent bias to on the regions selected. No animals were excluded from the statistical analysis.

## Supplementary Information

**Supplementary Figure 1:**
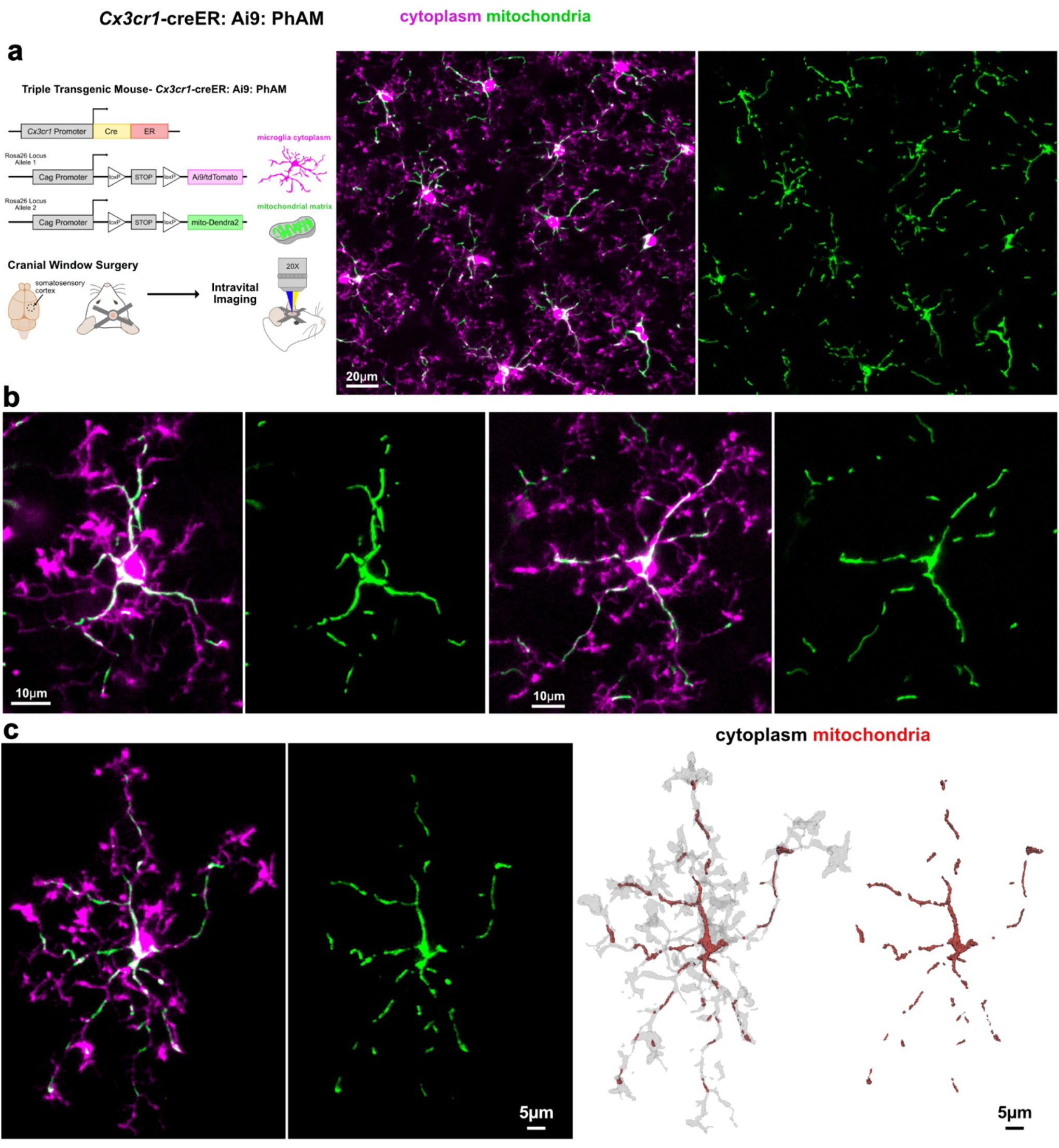
Mitochondrial labeling in microglia. **a)** Strategy for cre recombinase dependent conditional labeling of microglial cytoplasm and their mitochondria using floxed tdTomato and dendra2 fluorescent reporters. Microglia and their mitochondria were visualized in vivo using cranial window surgeries and intravital optical imaging. The image on the right shows an example of the labeling obtained in microglia of the superficial layers of the somatosensory cortex. **b)** Single microglia and their mitochondria. **c)** Fluorescent optical images and an example 3D reconstruction of a single microglia and its mitochondria.

**Supplementary Figure 2:**
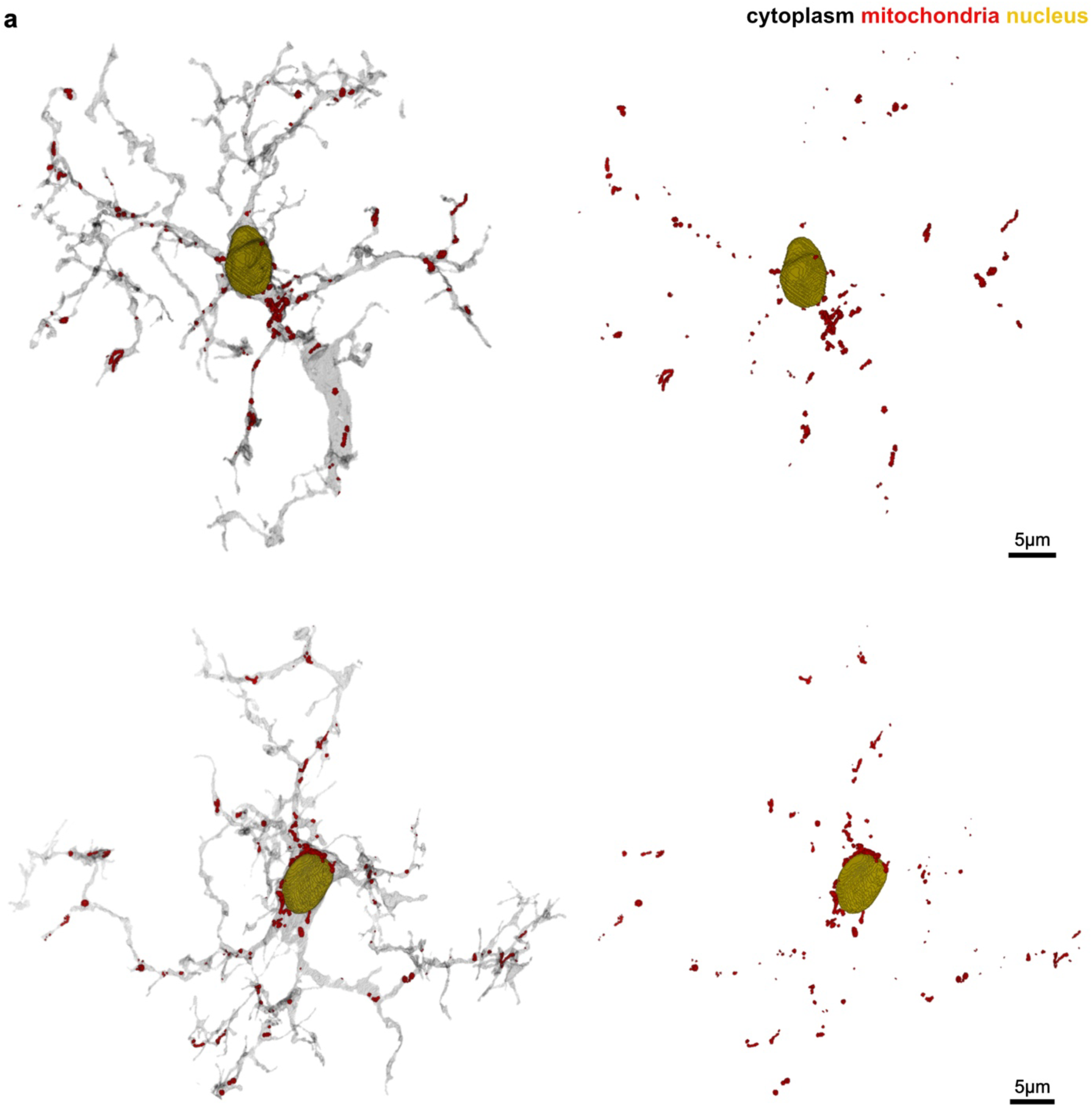
Mitochondria in human neocortical microglia. **a)** 3D reconstruction of two microglia (gray), their nuclei (yellow), and all their mitochondria (red) taken from a sample of human neocortex.

**Supplementary Figure 3:**
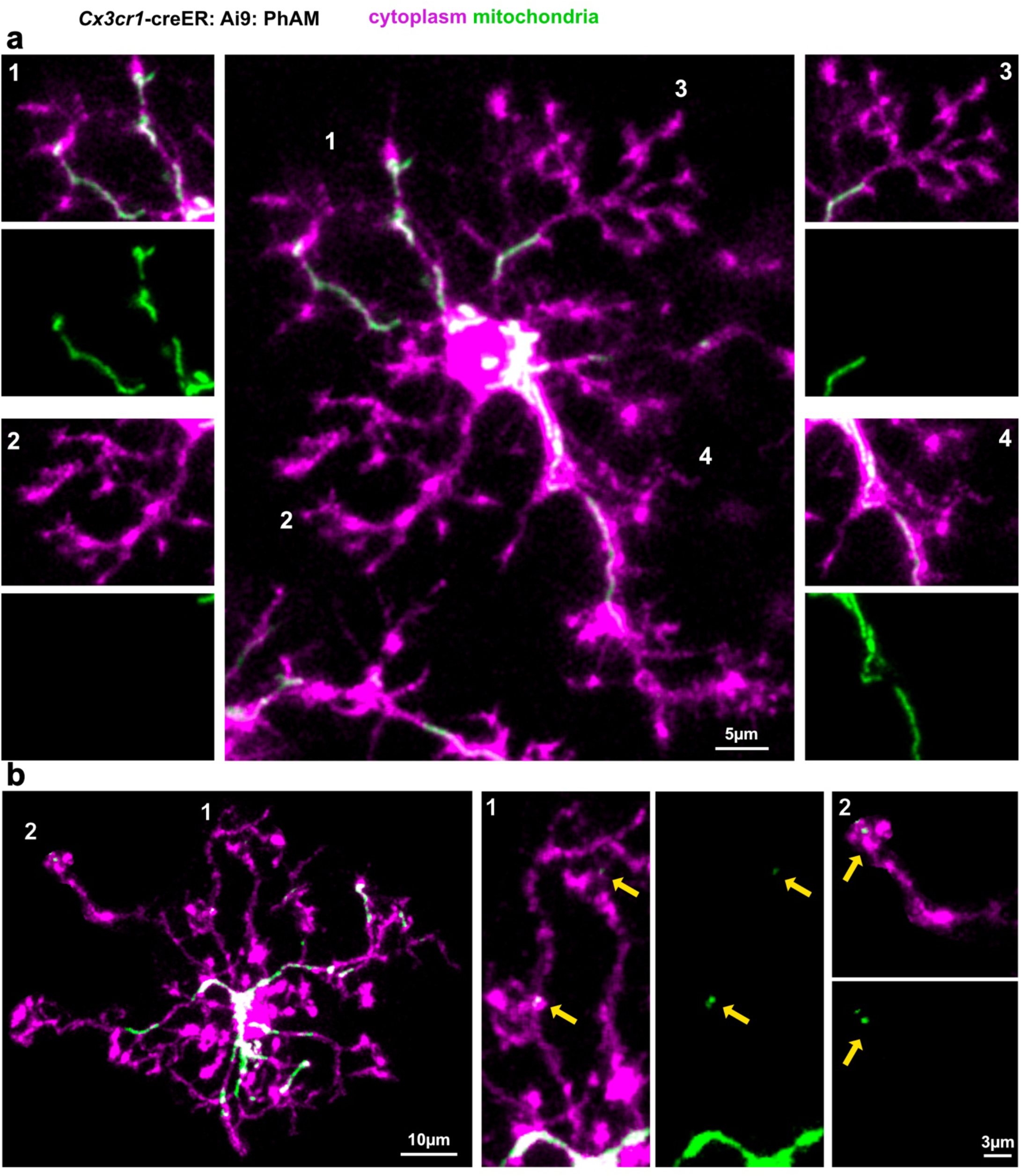
Heterogenous distribution of mitochondria in single microglial processes. **a)** In vivo image of a single microglia showing 4 different cell processes with varying densities of mitochondria showing the mitochondria are not uniformly distributed throughout the cell. **b)** Two different microglial processes with low mitochondrial volume that contain punctate, circular mitochondria (yellow arrows).

**Supplementary Figure 4:**
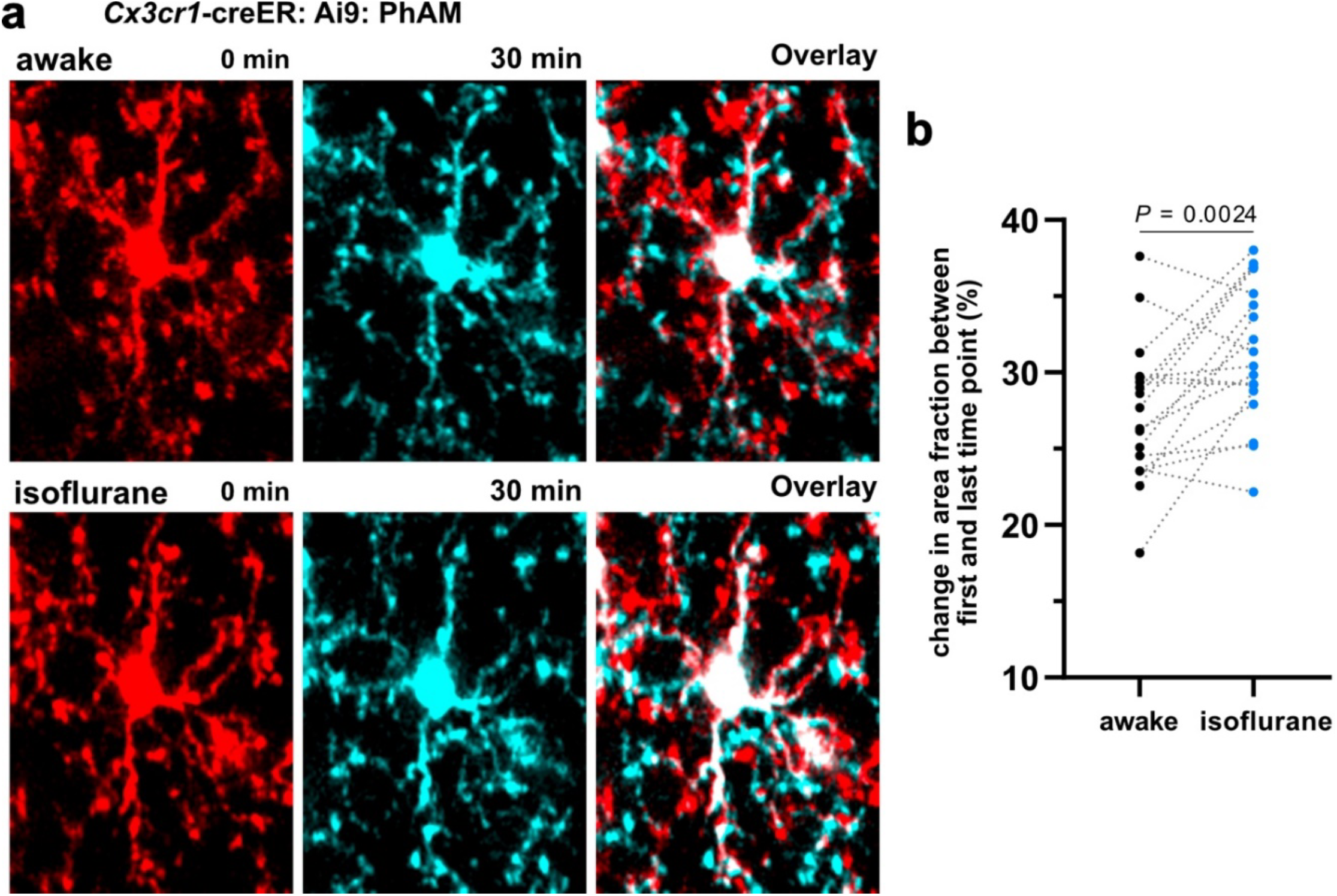
Microglial process surveillance is increased under anesthesia. **a)** Images showing microglia cytoplasm at 0 and 30 min taken from an awake (top) or isoflurane anesthetized (bottom) imaging session. Overlay of the two time points shows microglial process reorganization during the 30 minutes. **b)** Change in area fraction measurements of the two time points shown in (**a**) are used as a surrogate for measurements of microglial surveillance and the graph shows significantly more change in area fraction under the isoflurane condition compared to the awake state showing that microglial surveillance is higher under isoflurane. n = 18 cells from 6 mice; paired, two-tailed t-tests.

**Supplementary Figure 5:**
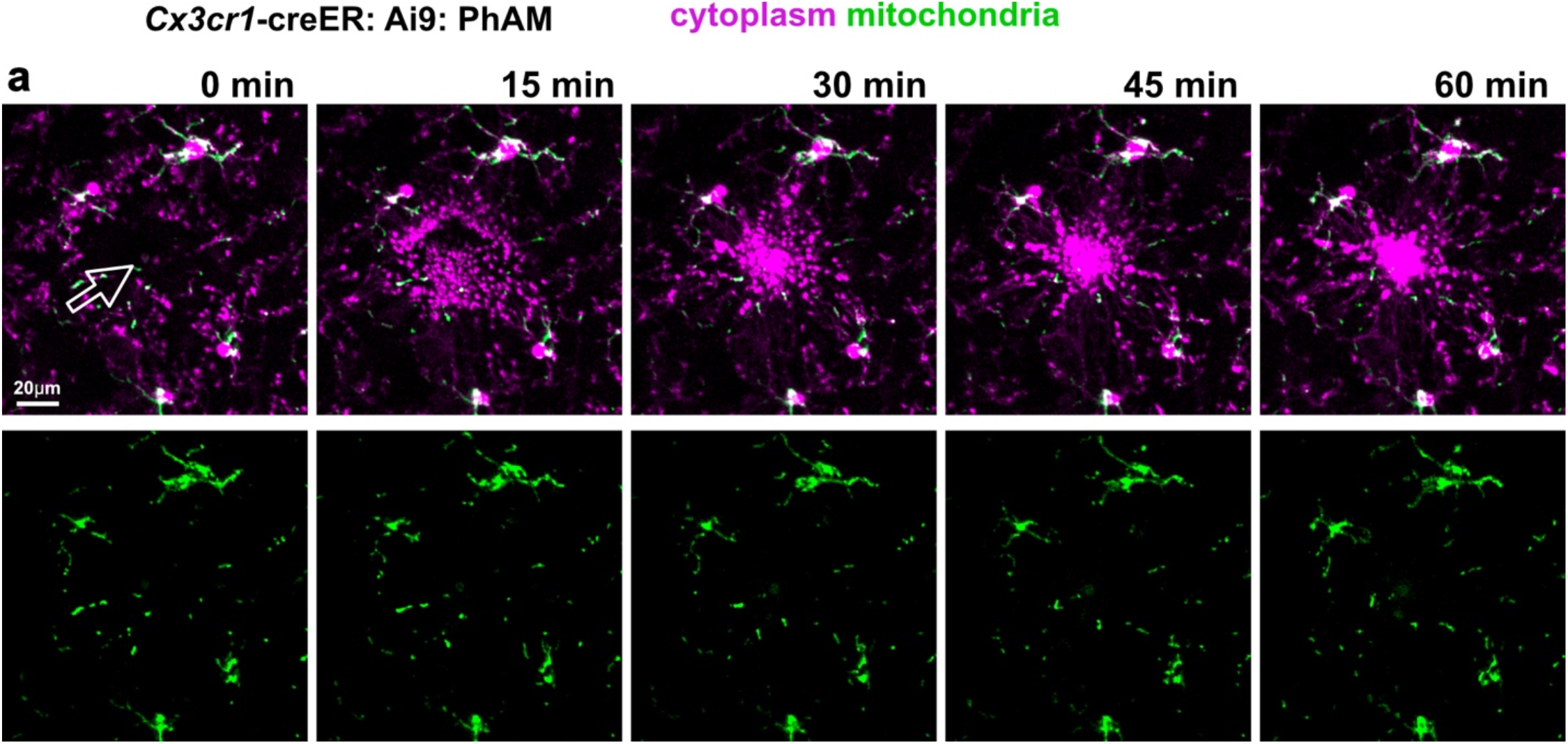
Microglial processes acutely responding to a laser lesion do not contain mitochondria. **a)** Time series showing the immediate chemotactic microglia process response to the site of a laser lesion (arrow), but little to no mitochondria response in the same time period.

**Supplementary Figure 6:**
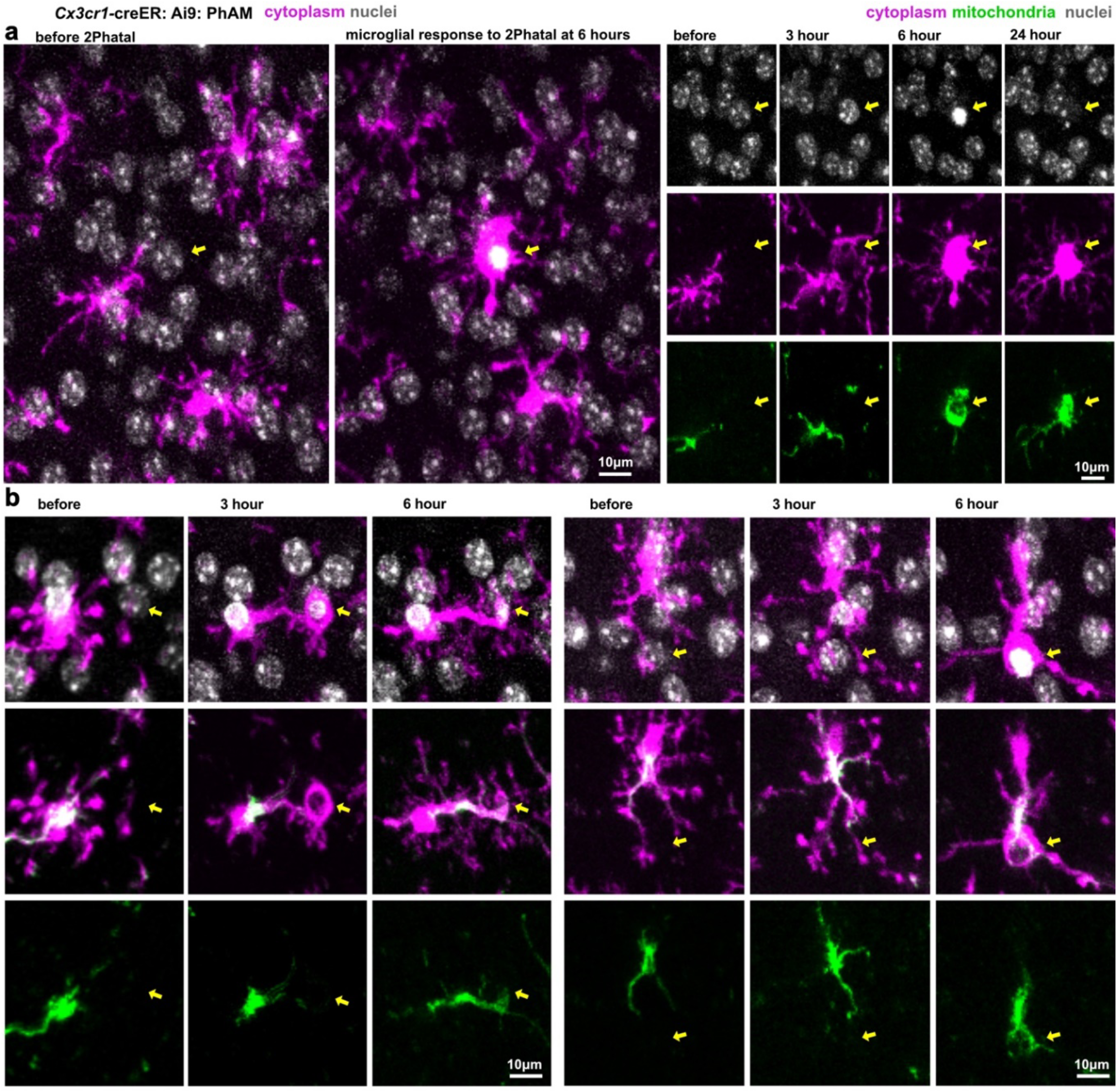
Mitochondria are delayed in their arrival to microglial processes engulfing dying neurons. **a)** Labeling of nuclei (gray) and microglia (magenta) indicating a cell that was targeted for 2Phatal (yellow arrows) before and 6 hours after the initiation of cell death. At 6 hours the microglial cell is engulfing the condensed nuclei of the targeted cell. The images on the right show the cytoplasm and mitochondria in the same microglia during the engulfment with very little mitochondrial presence at 3 hours during the initial stages of phagocytic engulfment. **b)** Two additional examples of phagocytic engulfment of dying neurons (yellow arrows) and the delayed presence of mitochondria in the phagocytic cup that forms around the condensed nuclei.

## Supplementary Movies

**Supplementary Movie 1: Mitochondrial labeling in mouse neocortical microglia**

Z-stack showing labeling of microglia and their mitochondria in the somatosensory cortex imaged through a cranial window. Depth from the cortical pial surface is shown in the top right corner.

**Supplementary Movie 2: Segmentation of mitochondria in human microglia**

3D rendering of a human microglial cell with its nucleus and all it its mitochondria segmented.

**Supplementary Movie 3: Intravital mitochondrial motility in neocortical microglia**

Time series of microglial surveillance and mitochondrial motility. The first clip in the movie shows multiple microglia and the subsequent clips focus in on single microglia and their mitochondria.

**Supplementary Movie 4: Mitochondrial motility in microglia is not altered by animal anesthesia**

Time series of a microglial cell and its mitochondria imaged in awake conditions and under isoflurane anesthesia.

**Supplementary Movie 5: Microglial processes responding to a laser lesion do not contain mitochondria**. Time series showing microglial process and mitochondrial chemotactic responses to focal laser lesions. The first video shows microglial motility for 30 minutes preceding the lesion followed by 45 min during the initial stages of process extension. The movie also shows a second example of a lesion and process extension towards the lesion in the subsequent 60 min with no mitochondria in the responding process.

## REFERENCES

1. Nimmerjahn, A., Kirchhoff, F. & Helmchen, F. Resting Microglial Cells Are Highly Dynamic Surveillants of Brain Parenchyma in Vivo. Science 308, 1314–1318 (2005).

2. Paolicelli, R. C. et al. Microglia states and nomenclature: A field at its crossroads. Neuron 110, 3458–3483 (2022).

3. Davalos, D. et al. ATP mediates rapid microglial response to local brain injury in vivo. Nat. Neurosci. 8, 752–758 (2005).

4. Hines, D. J., Hines, R. M., Mulligan, S. J. & Macvicar, B. A. Microglia processes block the spread of damage in the brain and require functional chloride channels. Glia 57, 1610–1618 (2009).

5. Hill, R. A., Damisah, E. C., Chen, F., Kwan, A. C. & Grutzendler, J. Targeted two-photon chemical apoptotic ablation of defined cell types in vivo. Nat. Commun. 8, 15837 (2017).

6. Damisah, E. C. et al. Astrocytes and microglia play orchestrated roles and respect phagocytic territories during neuronal corpse removal in vivo. Sci. Adv. 6, eaba3239 (2020).

7. Henson, P. M., Bratton, D. L. & Fadok, V. A. Apoptotic cell removal. Curr. Biol. 11, R795–R805 (2001).

8. Takahashi, K., Rochford, C. D. P. & Neumann, H. Clearance of apoptotic neurons without inflammation by microglial triggering receptor expressed on myeloid cells-2. J. Exp. Med. 201, 647–657 (2005).

9. Kawabori, M. & Yenari, M. A. The role of the microglia in acute CNS injury. Metab. Brain Dis. 30, 381–392 (2015).

10. Picard, M. & Shirihai, O. S. Mitochondrial signal transduction. Cell Metab. 34, 1620–1653 (2022).

11. Monzel, A. S., Enríquez, J. A. & Picard, M. Multifaceted mitochondria: moving mitochondrial science beyond function and dysfunction. Nat. Metab. 5, 546–562 (2023).

12. Liesa, M. & Shirihai, O. S. Mitochondrial Dynamics in the Regulation of Nutrient Utilization and Energy Expenditure. Cell Metab. 17, 491–506 (2013).

13. Fung, T. S., Chakrabarti, R. & Higgs, H. N. The multiple links between actin and mitochondria. Nat. Rev. Mol. Cell Biol. 1–17 (2023) doi:10.1038/s41580-023-00613-y.

14. Twig, G. et al. Biophysical properties of mitochondrial fusion events in pancreatic β-cells and cardiac cells unravel potential control mechanisms of its selectivity. Am. J. Physiol.-Cell Physiol. 299, C477–C487 (2010).

15. Marik, C., Felts, P. A., Bauer, J., Lassmann, H. & Smith, K. J. Lesion genesis in a subset of patients with multiple sclerosis: a role for innate immunity? Brain J. Neurol. 130, 2800–2815 (2007).

16. Streit, W. J., Braak, H., Xue, Q.-S. & Bechmann, I. Dystrophic (senescent) rather than activated microglial cells are associated with tau pathology and likely precede neurodegeneration in Alzheimer’s disease. Acta Neuropathol. (Berl.) 118, 475–485 (2009).

17. Damani, M. R. et al. Age-related alterations in the dynamic behavior of microglia. Aging Cell 10, 263–276 (2011).

18. Safaiyan, S. et al. Age-related myelin degradation burdens the clearance function of microglia during aging. Nat. Neurosci. 19, 995–998 (2016).

19. Mecca, C., Giambanco, I., Donato, R. & Arcuri, C. Microglia and Aging: The Role of the TREM2–DAP12 and CX3CL1-CX3CR1 Axes. Int. J. Mol. Sci. 19, 318 (2018).

20. Hill, R. A., Li, A. M. & Grutzendler, J. Lifelong cortical myelin plasticity and age-related degeneration in the live mammalian brain. Nat. Neurosci. 21, 683–695 (2018).

21. Angelova, D. M. & Brown, D. R. Microglia and the aging brain: are senescent microglia the key to neurodegeneration? J. Neurochem. 151, 676–688 (2019).

22. Leng, F. & Edison, P. Neuroinflammation and microglial activation in Alzheimer disease: where do we go from here? Nat. Rev. Neurol. 17, 157–172 (2021).

23. Choi, S., Guo, L. & Cordeiro, M. F. Retinal and Brain Microglia in Multiple Sclerosis and Neurodegeneration. Cells 10, 1507 (2021).

24. Malvaso, A. et al. Microglial Senescence and Activation in Healthy Aging and Alzheimer’s Disease: Systematic Review and Neuropathological Scoring. Cells 12, 2824 (2023).

25. McAvoy, K. & Kawamata, H. Glial mitochondrial function and dysfunction in health and neurodegeneration. Mol. Cell. Neurosci. 101, 103417 (2019).

26. Li, Y., Xia, X., Wang, Y. & Zheng, J. C. Mitochondrial dysfunction in microglia: a novel perspective for pathogenesis of Alzheimer’s disease. J. Neuroinflammation 19, 248 (2022).

27. Ulland, T. K. et al. TREM2 Maintains Microglial Metabolic Fitness in Alzheimer’s Disease. Cell 170, 649-663.e13 (2017).

28. Katoh, M. et al. Polymorphic regulation of mitochondrial fission and fusion modifies phenotypes of microglia in neuroinflammation. Sci. Rep. 7, 4942 (2017).

29. Nair, S. et al. Lipopolysaccharide-induced alteration of mitochondrial morphology induces a metabolic shift in microglia modulating the inflammatory response in vitro and in vivo. Glia 67, 1047–1061 (2019).

30. Montilla, A. et al. Role of Mitochondrial Dynamics in Microglial Activation and Metabolic Switch. ImmunoHorizons 5, 615–626 (2021).

31. Baik, S. H. et al. A Breakdown in Metabolic Reprogramming Causes Microglia Dysfunction in Alzheimer’s Disease. Cell Metab. 30, 493-507.e6 (2019).

32. Maes, M. E. et al. Mitochondrial network adaptations of microglia reveal sex-specific stress response after injury and UCP2 knockout. iScience 26, 107780 (2023).

33. Hu, Y. et al. Dual roles of hexokinase 2 in shaping microglial function by gating glycolytic flux and mitochondrial activity. Nat. Metab. 4, 1756–1774 (2022).

34. Stoolman, J. S. et al. Mitochondrial respiration in microglia is essential for response to demyelinating injury but not proliferation. Nat. Metab. 6, 1492–1504 (2024).

35. Mora-Romero, B. et al. Microglia mitochondrial complex I deficiency during development induces glial dysfunction and early lethality. Nat. Metab. 6, 1479–1491 (2024).

36. Peruzzotti-Jametti, L. et al. Mitochondrial complex I activity in microglia sustains neuroinflammation. Nature 628, 195–203 (2024).

37. Quintana-Cabrera, R. & Scorrano, L. Determinants and outcomes of mitochondrial dynamics. Mol. Cell 83, 857–876 (2023).

38. Shapson-Coe, A. et al. A petavoxel fragment of human cerebral cortex reconstructed at nanoscale resolution. Science 384, eadk4858 (2024).

39. Buchanan, J. et al. Oligodendrocyte precursor cells ingest axons in the mouse neocortex. Proc. Natl. Acad. Sci. 119, e2202580119 (2022).

40. Bernier, L.-P. et al. Nanoscale Surveillance of the Brain by Microglia via cAMP-Regulated Filopodia. Cell Rep. 27, 2895-2908.e4 (2019).

41. Stowell, R. D. et al. Noradrenergic signaling in the wakeful state inhibits microglial surveillance and synaptic plasticity in the mouse visual cortex. Nat. Neurosci. 22, 1782–1792 (2019).

42. Liu, Y. U. et al. Neuronal network activity controls microglial process surveillance in awake mice via norepinephrine signaling. Nat. Neurosci. 22, 1771–1781 (2019).

43. Misgeld, T. & Schwarz, T. L. Mitostasis in Neurons: Maintaining Mitochondria in an Extended Cellular Architecture. Neuron 96, 651–666 (2017).

44. Faitg, J. et al. 3D neuronal mitochondrial morphology in axons, dendrites, and somata of the aging mouse hippocampus. Cell Rep. 36, (2021).

45. Jackson, J. G. & Robinson, M. B. Regulation of mitochondrial dynamics in astrocytes: mechanisms, consequences, and unknowns. Glia 66, 1213–1234 (2018).

46. Hill, R. A., Nishiyama, A. & Hughes, E. G. Features, Fates, and Functions of Oligodendrocyte Precursor Cells. Cold Spring Harb. Perspect. Biol. 16, a041425 (2024).

47. Bame, X. & Hill, R. A. Mitochondrial network reorganization and transient expansion during oligodendrocyte generation. Nat. Commun. 15, 6979 (2024).

48. Han, S. M., Baig, H. S. & Hammarlund, M. Mitochondria Localize to Injured Axons to Support Regeneration. Neuron 92, 1308–1323 (2016).

49. Cunniff, B., McKenzie, A. J., Heintz, N. H. & Howe, A. K. AMPK activity regulates trafficking of mitochondria to the leading edge during cell migration and matrix invasion. Mol. Biol. Cell 27, 2662–2674 (2016).

50. Desai, S. P., Bhatia, S. N., Toner, M. & Irimia, D. Mitochondrial Localization and the Persistent Migration of Epithelial Cancer cells. Biophys. J. 104, 2077–2088 (2013).

51. Crosas-Molist, E. et al. AMPK is a mechano-metabolic sensor linking cell adhesion and mitochondrial dynamics to Myosin-dependent cell migration. Nat. Commun. 14, 2740 (2023).

52. Shneyer, B. I., Ušaj, M., Wiesel-Motiuk, N., Regev, R. & Henn, A. ROS induced distribution of mitochondria to filopodia by Myo19 depends on a class specific tryptophan in the motor domain. Sci. Rep. 7, 11577 (2017).

53. Uhlemann, R. et al. Actin dynamics shape microglia effector functions. Brain Struct. Funct. 221, 2717–2734 (2016).

54. Möller, K. et al. A role for the centrosome in regulating the rate of neuronal efferocytosis by microglia in vivo. eLife 11, e82094 (2022).

55. Kim, S. Y. et al. Neuronal mitochondrial morphology is significantly affected by both fixative and oxygen level during perfusion. Front. Mol. Neurosci. 15, 1042616 (2022).

56. Kelly, B. & O’Neill, L. A. Metabolic reprogramming in macrophages and dendritic cells in innate immunity. Cell Res. 25, 771–784 (2015).

57. Voloboueva, L. A., Emery, J. F., Sun, X. & Giffard, R. G. Inflammatory response of microglial BV-2 cells includes a glycolytic shift and is modulated by mitochondrial glucose-regulated protein 75/mortalin. FEBS Lett. 587, 756–762 (2013).

58. Rabbani, N. & Thornalley, P. J. Hexokinase-2 Glycolytic Overload in Diabetes and Ischemia–Reperfusion Injury. Trends Endocrinol. Metab. 30, 419–431 (2019).

59. Paolicelli, R. C. & Pluchino, S. Complex roles for mitochondrial complexes in microglia. Nat. Metab. 6, 1426–1428 (2024).

60. He, D. et al. Disruption of the IL-33-ST2-AKT signaling axis impairs neurodevelopment by inhibiting microglial metabolic adaptation and phagocytic function. Immunity 55, 159-173.e9 (2022).

61. Yona, S. et al. Fate Mapping Reveals Origins and Dynamics of Monocytes and Tissue Macrophages under Homeostasis. Immunity 38, 79–91 (2013).

62. Madisen, L. et al. A robust and high-throughput Cre reporting and characterization system for the whole mouse brain. Nat. Neurosci. 13, 133–140 (2010).

63. Pham, A. H., McCaffery, J. M. & Chan, D. C. Mouse lines with photo-activatable mitochondria to study mitochondrial dynamics. genesis 50, 833–843 (2012).

64. Jackson, S. J. et al. Does age matter? The impact of rodent age on study outcomes. Lab. Anim. 51, 160–169 (2017).

65. Chapman, T. W., Olveda, G. E., Bame, X., Pereira, E. & Hill, R. A. Oligodendrocyte death initiates synchronous remyelination to restore cortical myelin patterns in mice. Nat. Neurosci. 1–15 (2023) doi:10.1038/s41593-023-01271-1.

66. Olveda, G. E., Barasa, M. N. & Hill, R. A. Microglial phagocytosis of single dying oligodendrocytes is mediated by CX3CR1 but not MERTK. Cell Rep. 43, (2024).

67. Berger, D. R., Seung, H. S. & Lichtman, J. W. VAST (Volume Annotation and Segmentation Tool): Efficient Manual and Semi-Automatic Labeling of Large 3D Image Stacks. Front. Neural Circuits 12, (2018).

68. Fiji Downloads. ImageJ Wiki https://imagej.github.io/software/fiji/downloads.

69. MATLAB version: 24.1.0 (R2024a). Natick, Massachusetts: The MathWorks Inc.; 2024 https://www.mathworks.com.

70. 3D Slicer image computing platform. 3D Slicer https://slicer.org/.

71. Kikinis, R., Pieper, S. D. & Vosburgh, K. G. 3D Slicer: A Platform for Subject-Specific Image Analysis, Visualization, and Clinical Support. in Intraoperative Imaging and Image-Guided Therapy (ed. Jolesz, F. A.) 277–289 (Springer, New York, NY, 2014). doi:10.1007/978-1-4614-7657-3_19.

72. Kapur, T. et al. Increasing the impact of medical image computing using community-based open-access hackathons: The NA-MIC and 3D Slicer experience. Med. Image Anal. 33, 176–180 (2016).

73. Fedorov, A. et al. 3D Slicer as an Image Computing Platform for the Quantitative Imaging Network. Magn. Reson. Imaging 30, 1323–1341 (2012).

74. Ershov, D. et al. TrackMate 7: integrating state-of-the-art segmentation algorithms into tracking pipelines. Nat. Methods 19, 829–832 (2022).

75. Ohsawa, K. et al. P2Y12 receptor-mediated integrin-beta1 activation regulates microglial process extension induced by ATP. Glia 58, 790–801 (2010).

